# Long-Term Intra-Host Evolution of SARS-CoV-2 in an Immunocompromised Patient: Recombination and Within-Host Mutations Driving Viral Adaptation

**DOI:** 10.1101/2025.03.25.645210

**Authors:** Emilie Burel, Pierre Pontarotti, Jacques Fantini, Jean-Christophe Lagier, Pierre Edouard Fournier

**Affiliations:** IHU Méditerranée Infection, 19-21 boulevard Jean Moulin, 13005 Marseille, France; RITMES, Aix-Marseille Univ., Service de Santé des Armées, IHU Méditerranée Infection, Marseille, France; MEPHI, Aix-Marseille Univ., IHU Méditerranée Infection, Marseille, France; INSERM UMR UA 16, Aix-Marseille Université, Marseille, France; Pôle de maladies infectieuses, Assistance Publique - Hôpitaux de Marseille (AP-HM), IHU Méditerranée Infection, Marseille, France; CNRS SNC5039, Marseille, France

**Author notes:** **Corresponding author:** Pierre Edouard Fournier, IHU - Méditerranée Infection, 19-21 boulevard Jean Moulin, 13005 Marseille, France.

**Keywords:** Intra-host evolution, SARS-CoV-2, recombination, genomics, iSNV, Spike

## Abstract

An immuno-compromised patient with lymphoma experienced a prolonged SARS-CoV-2 infection lasting 14 months, initially infected with B.1.160, followed by B.1.1.7. This study focused on intra-host single nucleotide variants (iSNVs) and single nucleotide polymorphisms (SNPs), distinguishing their origin as either within-host mutations or parental. The whole genome analysis revealed accelerated evolution, with positive selection detected in key genes such as Spike, N, ORF9b, and nsp13, all involved in viral replication and immune evasion. Of the two evolutionary mechanisms involved, host-driven mutation has played a dominant role in this evolutionary story. C>T transitions emerged as the most widespread mutational signature, consistent with host-driven RNA editing mechanisms. Two host-internal mutations, A28271T in the translation initiation region of the N Kozak gene and C26858T in the M gene, were highly shared among the deposited SARS-CoV-2 sequences. Recombination with parental lineages played a major role particularly in Spike, ORF3a and M genes. In Spike, the B.1.1.7 sequence was selected, wild-type ORF3a and M were restored, suggesting a selective advantage in returning to an ancestral sequence. In addition, the temporary emergence of intra-host convergent mutations in Spike, notably L5F, D796H and T572I, underlines the strong selective pressures exerted on this gene. A 126-nucleotide deletion in ORF8 resulted in a truncated protein, reinforcing its uselessness for viral replication, as observed in circulating variants such as B.1.1.7. The present case highlights the complex interplay between viral recombination and mutations within the host in chronic infections and further underscores the remarkable evolutionary plasticity of SARS-CoV-2 and its potential to generate highly adapted viral strains.

## INTRODUCTION

The COVID-19 pandemic has been shaped by the regular emergence of SARS-CoV-2 variants with properties that escape natural or vaccine-induced immunity, prompting interest in understanding their origins. Intra-host evolution in long-term infections has emerged as a possible origin^1,2^, a theory supported in particular by the emergence of the Alpha (B.1.1.7)^3^ and Omicron (B.1.1.529) variants^4^, which quickly acquired numerous mutations, in comparison with variants circulating during the previous epidemic waves, conferring immune escape and increased transmissibility. Documenting intra-host evolution that leads to unique patterns is crucial for understanding the mechanisms driving the emergence of host-adapted strains, such as those with increased affinity for the ACE2 Spike receptor^5,6^. Immunocompromised patients, due to AIDS, some cancers or immunosuppressive therapy, for the main cases, are more prone to develop long-term infections due to impaired viral clearance^7,8,9,10,11,12–15^. In addition, weakened immune pressure, combined with treatments such as antiviral therapies, antibodies and convalescent plasma, create favorable conditions for the emergence of new genotypes^16,10,5,17,18^ and convergent events^19^. During prolonged infections, co-infections with multiple strains have been documented^20,21^ which can lead to recombination through intracellular co-infection and RNA polymerase-induced homologous recombination^22,23^. It has been suggested that this process played a key role in the emergence of SARS-CoV-2^24,25^, being particularly impactful on the Spike protein gene^26,27,28,29^. Recombination was extensively described for Omicron variants and subvariants^30,31^ and it has recently been shown that SARS-CoV-2 has a higher frequency of RNA recombination than other human CoVs^32^. Moreover, the study of intra-host nucleotide variation (iSNV) is crucial for understanding the early mutagenic mechanisms at the origin of the creation of a pool of quasispecies^33,34,35^. Among the dominant mutations observed during intra-host evolution of SARS-CoV-2, C>T transitions are particularly frequent, and have been linked to the activity of enzymes from the APOBEC (Apolipoprotein B mRNA Editing Catalytic Polypeptide-like) family, including APOBEC3A, APOBEC1, and APOBEC3G, which are cytidine deaminases^36,37^. Combined mechanisms involving host-induced RNA editing and host environmental pressure associated with recombination, may thus contribute to the rapid evolution of viruses.

We previously described and confirmed by culture the emergence of recombinant strains in a patient infected with B.1.160, who was later superinfected by B.1.1.7, resulting in unique strains^38^. Given the exceptional duration of this chronic infection (440 days), this case provided a rare opportunity to investigate intra-host viral evolution over an extended period which combined both mechanisms: recombination and within host emergence of mutations, offering valuable insights into their interplay. To further explore this evolutionary process, we conducted ultra-deep sequencing on 16 nasopharyngeal swabs collected over time using Illumina technology for all samples. Twelve samples offered sufficient coverage in order to make a deep analysis on iSNVs and SNPs patterns.

Our findings revealed extensive intra-host diversity, with positive selection observed in key genes such as N, ORF9b, nsp4, nsp13 and nsp3, highlighting their adaptive potential. Additionally, we identified a convergent event in (A28271T) substitution in a non-coding region, located in the Kozak sequence upstream of the Nucleocapsid gene, potentially optimizing viral gene expression^39^. In the Spike gene, we observed transient convergent mutations: D796H and L5F, which were subsequently eliminated by recombination, while within-host mutations T572I and T618K appeared among the parental B.1.1.7 haplotype. Moreover, recombination led to the restoration of the wild-type (WT) ORF3a and WT-like M sequence, suggesting a selective advantage for reverting to the ancestral sequence. This event emphasizes the role of recombination not only in generating genetic diversity but also in purging mutations. Furthermore, we identified a 126-nt deletion in ORF8, resulting in a truncated protein form, reinforcing its dispensability for viral replication, as previously observed in circulating variants such as B.1.1.7.

Finally, this case provides a unique example of intra-host evolution of SARS-CoV-2, testifying to its genomic plasticity generating highly host-adapted genotypes, notably an optimized spike, and also evidenced by recombination reversions and the significant accumulation of new within host mutations. The interplay between host-induced mutagenesis, selection pressures and recombination provides a rich framework for study, which could be used in future comparative studies aimed at better understanding host adaptation strategies.

## MATERIALS AND METHODS

### Ethics statement

This retrospective study received a positive decision from the “Assistance Publique-Hôpitaux de Marseille”(AP-HM) ethics and scientific committee (N° CSE24-4). Access to biological and patient registry data from the hospital information system was approved by the AP-HM data protection committee and registered in the European General Data Protection Regulation register under number RGPD/APHM 2019-73.

### Samples

This study was carried out retrospectively on 54 nasopharyngeal swabs collected from September 2020 to December 2021, as part of care of one patient at the IHU Mediterranee Infection Institute (Marseille, France).

### SARS-CoV-2 Genome Sequencing

Illumina sequencing was performed only from samples with Ct values <30, as previously described^38^. Viral RNA was extracted from 200 μL of nasopharyngeal swab fluid using the EZ1 Virus Mini kit v2.0 on an EZ1 Advanced XL instrument (Qiagen, Courtaboeuf, France) or using the MagMax Viral/Pathogen Nucleic Acid Isolation kit on the KingFisher Flex system (Thermo Fisher Scientific, Waltham, MA, USA), following the manufacturer’s instructions.

Next-generation sequencing performed on the NovaSeq 6000 instrument followed the Illumina COVIDSeq protocol (Illumina Inc; Illumina Inc., San Diego, CA, USA.), which included first strand cDNA synthesis from extracted viral RNA; cDNA amplification with two COVIDSeq primer pools; fragmentation and tagging of PCR amplicons with adapter sequences; clean up; PCR amplification with index ligation (seven cycles) of tagmented amplicons; pool, clean up, quantification and normalization of libraries; library sequencing on a NovaSeq 6000 sequencing system SP flow cell (Illumina Inc; Illumina Inc., San Diego, CA, USA).

### Complete genome analysis

Raw files were basecalled using the Dragen Bcl Convert pipeline (v3.9.3; https://emea.support.illumina.com/sequencing/sequencing_software/bcl-convert.html; accessed on 18 May 2022), generating fastq files. Raw reads were trimmed using Trimmomatic (V 0.39) (https://doi.org/10.1093/bioinformatics/btu170) and then mapped using bwa-mem2 (Sequence alignment using Burrows-Wheeler Transform) (v2.2.1; https://github.com/bwa-mem2/bwa-mem2; 2019 Intel Corporation, Heng Li) against the Wuhan-Hu-1 isolate reference (GenBank NC_045512.2); primers were cleaned using Samtools ampliconclip (v1.13; https://www.htslib.org/; accessed on 18 May 2022). Variant calling was performed using FreeBayes (v1.3.5; https://github.com/freebayes/freebayes; accessed on 18 May 2022, Erik Garrison, Gabor Marth, University of Tennessee and Utah) and consensus genomes were obtained with Bcftools (v1.13; https://samtools.github.io/bcftools/bcftools.html; accessed on 18 May 2022, 2012–2021 Genome Research Ltd.). The consensus threshold of a SNP (Single Nucleotide Polymorphism) has been set at 70% with a minimum coverage of 10 reads.

To check for the presence of large Indels, given the length of the reads (50 bp), the cleaned reads were also assembled with Unicycler (v0.4.4) (https://doi.org/10.1371/journal.pcbi.1005595), three almost complete genomes were obtained in one contig, the resulting assemblies were deposited in the GISAID database (https://gisaid.org/) (EPI_ISL_18896020, EPI_ISL_18897181, EPI_ISL_18897180).

RNA sequences in this study are represented using thymine (T) instead of uracil (U), in order to facilitate sequence comparisons and computational processing. It should obviously be understood that T represents U in its biological context.

Substitutions detected after a deletion (ex : G28038T) were excluded from the analysis because of artifactual origin.

### In depth analysis of intra host diversity

To investigate intra-host SARS-CoV-2 evolution and diversity, we classified mutations based on their origin and allele frequency (AF). Within-host mutations were defined as novel mutations absent from both parental lineages (B.1.160 and B.1.1.7) and were obtained by excluding all signature mutations from B.1.160 and B.1.1.7. Although full reconstruction of the original B.1.1.7 haplotype was not possible due to lack of pre-D229 samples, a detailed analysis of contemporaneous B.1.1.7 strains in Marseille found no evidence of externally introduced mutations (Table S1). Therefore, all mutations except those defining B.1.160 or B.1.1.7 were considered to be internal host mutations, and additional external genomic contributions negligible.

To ensure high-confidence variant detection, we set a minimum allele frequency (MAF) threshold of 5% and a minimum sequencing depth of 50x, following recommendations from previous studies^40^. To further refine variant calling, only iSNVs covered by at least 5 reads were retained, ensuring a practical MAF threshold of ≥10% at 50x depth. Additionally, a genome-wide coverage threshold of >80% was applied to ensure the inclusion of high-quality samples in the analysis at 50x coverage.

iSNV analysis was performed on BAM files obtained from the previously described mapping process. Variant calling was conducted using iVar (v1.3.1, https://github.com/andersen-lab/ivar) in combination with Samtools (https://github.com/samtools/samtools), with the precited parameters and a minimum Phred quality score of 30.

VCF files generated by iVar and TSV files from the mapping step were processed using custom Python scripts (https://github.com/EMbBureli/Intra-host-SARS-CoV-2-evolution-study-in-an-immunocompromised-patient) to generate filtered mutation files.

### Phylogenetic analysis

The phylogenetic analysis was obtained with the Nexstrain pipeline^41^ using the Maximum-likelihood method^42^. The intra-host evolutionary rate was calculated and compared to inter-host rates from 700 locally circulating SARS-CoV-2 sequences (GISAID dataset, September 2020–December 2021). Plots were visualized using the Auspice online tool (https://auspice.us/).

### Entropy calculation based at gene level

Shannon entropy (H) was used to measure the mutational diversity of allele frequencies at gene levels within each pool and was computed using the following formula:

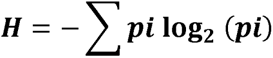

where pi represents the frequency of alternative alleles for a given mutation. To minimize bias due to variable gene coverage, the raw entropy values were divided by the percentage of gene coverage. The normalized entropy values per gene were summed to obtain an overall measure of mutation diversity, and pairwise statistical significance was applied using the Mann-Whitney U test, from scipy.stats (https://docs.scipy.org/doc/scipy/) Python library. Graphs were plotted using the Matplotlib Python library (https://matplotlib.org/stable/plot_types/).

### pN/pS calculation

The pN/pS ratio was computed following the Nei-Gojobori method^43^, without the Jukes-Cantor correction for multiple substitutions, given the relatively short evolutionary time within the host, with 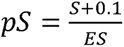 and 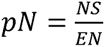, where S and NS represent the number of observed synonymous and non-synonymous mutations respectively, within a given gene. The corresponding synonymous (ES) and non-synonymous (EN) sites were calculated using a Python script based on the reference SARS-CoV-2 genome (NC_045512.2). For cases where pS = 0 and pN =0, the impossible ratio was annotated as “undetermined”, and “infinite” when NS>0. The computed pN/pS ratio, number of infinite and undetermined cases were plotted using the python Matplotlib library (https://matplotlib.org/stable/plot_types/).

### Molecular modeling of mutant SARS-CoV-2 Spike proteins

Molecular modeling of mutant spike proteins was performed as described previously^44^. A complete structure of the original SARS-CoV-2 spike protein encompassing the amino acid residues 14-1200 (thus lacking the signal sequence) was generated and characterized as described^45^. Minimized structures of the spike protein of each variant were obtained by sequentially introducing appropriate mutations and/or deletions. Energy minimizations of the variants were performed with the Polak-Ribière conjugate gradient algorithm using a maximum of 3 × 105 steps and a root mean square (RMS) gradient of 0.01 kcal/mol.Å as the convergence condition^46^. The electrostatic potential illustrated in this study is the sum of the Coulomb potentials for each atom of the considered molecule, with a distance-dependent dielectric constant^47^.

### Within-Host Mutations and Global Convergence

To assess the prevalence of SNPs detected within-host (SNP_WH) in globally circulating SARS-CoV-2 strains, we queried the GISAID (Global Initiative on Sharing Avian Influenza Data, https://www.gisaid.org) database on November 30, 2023, analyzing a dataset of 16,269,440 SARS-CoV-2 genomes. The visualization of SNP_WH proportions in the global dataset was generated using the ggplot2 library in R (https://ggplot2.tidyverse.org/).

### Mapping visualization

The ORF8 deletion was directly visualized from PCR products realized on D378 sample targeting region 24,813-29,074. Reads were mapped against the SARS-CoV-2 reference genome. Sam files were compressed into .bam files with Samtools, (v1.13; https://www.htslib.org/), indexed and visualized using the IGV software (https://www.igv.org/)^48^, the 126-nt deletion was visible thanks to continuous reads.

All scripts are available at :(https://github.com/EMbBureli/Intra-host-SARS-CoV-2-evolution-study-in-an-immunocompromised-patient).

## RESULTS

### An immunocompromised patient with a chronic SARS-CoV-2 infection

A 56-year-old immunocompromised patient with a severe pneumonia was diagnosed with SARS-CoV-2 infection in September 2020, resulting in admission to intensive care. Despite the improvement in his condition, he remained SARS-CoV-2-positive for 14 months (Figure 1a-b). In addition, in May 2021, a progressive multifocal leukoencephalitis was diagnosed. In his medical history, the patient was previously diagnosed with mixed Hodgkin and follicular lymphoma in 2017. The patient was still under therapy for the follicular lymphoma in 2021. In terms of infecting genotype, the patient first contracted the SARS-CoV-2 variant B.1.160 in August 2020 and then B.1.1.7, which was partially detected from May 2021 (Figure 1 a-d), based on 19 lineage-defining mutations. The evolutionary process involved multiple recombination events and the appearance of within host mutations. Different recombinant regions from the B.1.1.7 variant were identified during the infection (Figure 1d) and objectified in the previous article^38^, including the 5’ end, comprising C913T and C3267T, positions 17,019 to 26,876, including the Spike gene (21,563-25,384) at consensus level, and positions 27,972 to 28,977 at sub consensus level. Notably, the sample corresponding to day 229 contained 11 additional B.1.1.7 lineage-defining mutations with allele frequencies (AF) ranging between 13.3% and 36.4%, reflecting early stages of recombination before these mutations were eventually eliminated (Figure S2). Treatment regimens received by the patient included two convalescent plasma treatments as well as Lenalidomide, Rituximab and Pembrolizumab chemotherapies. The serological data for May and October 2021 are presented in Table 1. The patient died in December 2021 from complications of his hematological and neurological diseases.

**Figure 1:**
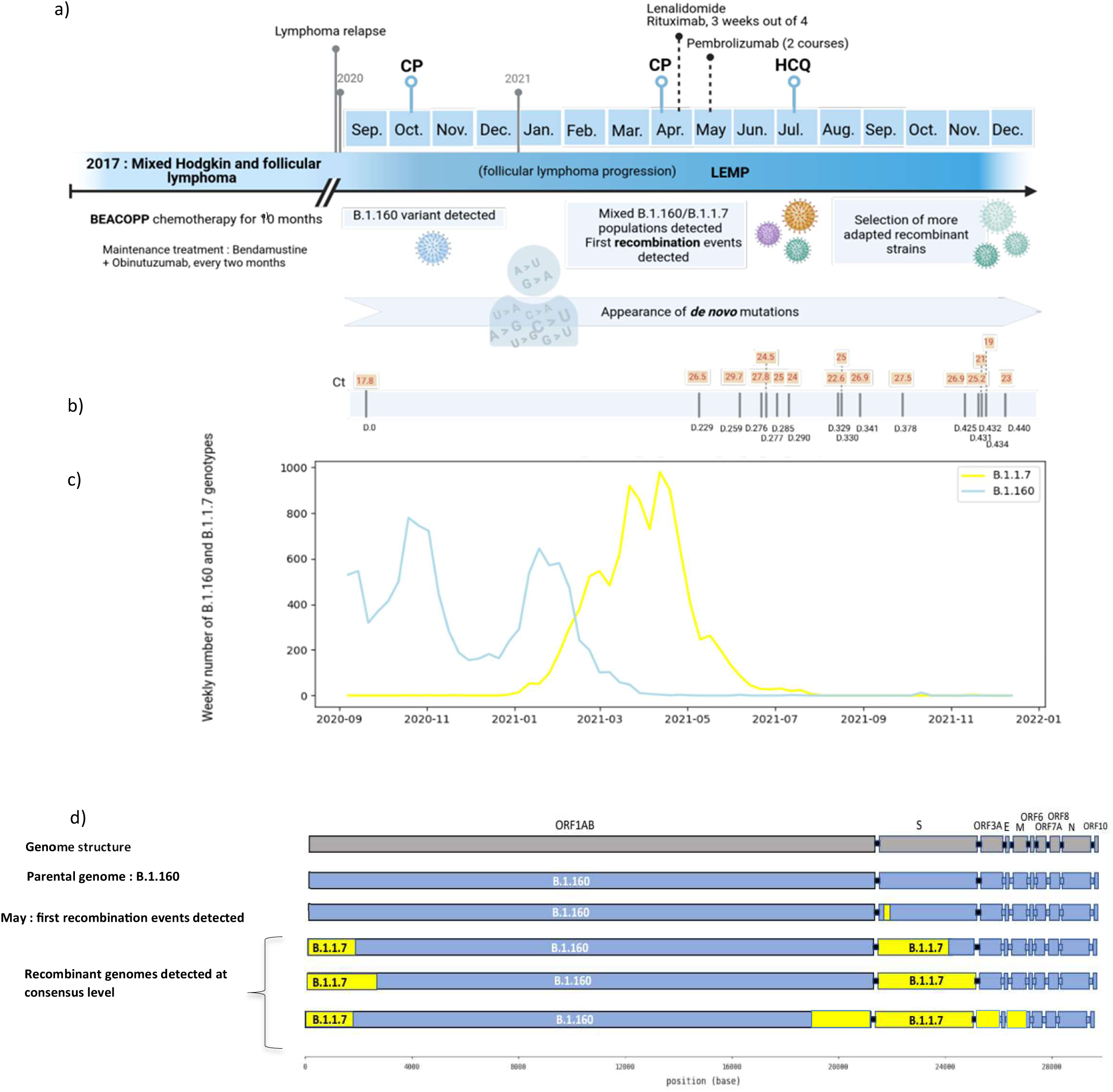
Timeline of the patient care and SARS-CoV-2 variant evolution undergoing recombination and *de novo* emergence of mutations. a) Timeline of the patient’s treatment regimen, including chemotherapy, immunotherapy, and antiviral interventions, in relation to the detection of SARS-CoV-2 variants. b) The sequenced nasopharyngeal samples are marked on the timeline with their corresponding qPCR Ct values. c) Superimposition of the epidemiological curves of the two parental genomes, B.1.160 and B.1.1.7 lineages, that infected the patient, in Marseille, the patient’s city of residence. d) Genomic structure of the detected recombinant consensus-level strains. The parental B.1.160 genome is represented in blue, while segments originating from the B.1.1.7 variant are shown in yellow, illustrating the progressive acquisition of recombination-derived genomic regions.

**Table 1:** Serological data obtained on serum.

| Date | Nature | Technical | Results : IgG anti SARS-CoV-2 |  |
| --- | --- | --- | --- | --- |
| 2021-05-03 | Serum | Immunoluminometric technique Binding | Ti Index 16 | Positive |
| 2021-05-25 | Serum | Immunoluminometric technique Binding | Ti Index 10 | Negative |
| 2021-08-12 | Serum | Immunoluminometric technique Binding | Ti Index <4 | Negative |
| 2021-11-23 | Serum | Immunoluminometric technique Binding | Ti Index <5 | Negative |

### Genomic evolution of SARS-CoV-2 over time, a global overview

As part of the longitudinal monitoring of the virus’ evolution, we first focused on the analysis of 16 consensus genomes, which are artificial reconstructions based on the dominant nucleotide at each position, obtained from nasopharyngeal swabs taken between September 2020 and December 2021. These genomes were sequenced from samples with Ct values below 30, all sequences were classified as B.1.160 variants according to the PangoLIN (Phylogenetic Assignment of Named Global Outbreak LINeages) nomenclature (https://cov-lineages.org/). The GISAID accession numbers for these sequences are presented in Table 2. Phylogenetic reconstruction based on the 16 consensus genomes (≥10× coverage) revealed the emergence of three distinct genotypes over time (Figure 2a): Genotype 1 was dominant at days 229, 259, and 276; Genotype 2 was detected at days 277, 290, 329, 341 and 378; and Genotype 3 emerged at days 285, 330, 431, 432, 425, 434, and 440.

**Table 2:** Summary of SARS-CoV-2 genomes used in this study. Table showing sequencing information for the 16 SARS-CoV-2 consensus genomes obtained with a 10x coverage. Displayed are each sample’s GISAID identifier, strain name (associated with sampling date) and PangoLIN lineage.

| GISAID Identifier | Strain name (Time for Diagnosis) | Sampling date | PangoLIN lineage | Coverage | NGS Technology, Instrument |
| --- | --- | --- | --- | --- | --- |
| EPI_ISL_6332079 | D0 | 2020-09-18 | B.1.160 | 99.30 % | Illumina, NovaSeq |
| EPI_ISL_10816743 | D229 | 2021-05-05 | B.1.160 | 97.10 % | Illumina, NovaSeq |
| EPI_ISL_19195977 | D259 | 2021-06-04 | B.1.160 | 91.80 % | Illumina, NovaSeq |
| EPI_ISL_11030507 | D276 | 2021-06-21 | B.1.160 | 90.20 % | Illumina, NovaSeq |
| EPI_ISL_19195978 | D277 | 2021-06-22 | B.1.160 | 96.80 % | Illumina, NovaSeq |
| EPI_ISL_19195979 | D285 | 2021-06-30 | B.1.160 | 97.90 % | Illumina, NovaSeq |
| EPI_ISL_10816731 | D290 | 2021-07-05 | B.1.160 | 97.70 % | Illumina, NovaSeq |
| EPI_ISL_19195980 | D329 | 2021-08-13 | B.1.160 | 99.20 % | Illumina, NovaSeq |
| EPI_ISL_19195981 | D330 | 2021-08-14 | B.1.160 | 97.40 % | Illumina, NovaSeq |
| EPI_ISL_19195982 | D341 | 2021-08-25 | B.1.160 | 83.60 % | Illumina, NovaSeq |
| EPI_ISL_10816744 | D378 | 2021-10-01 | B.1.160 | 95.80 % | Illumina, NovaSeq |
| EPI_ISL_19196018 | D425 | 2021-11-17 | B.1.160 | 82.60 % | Illumina, NovaSeq |
| EPI_ISL_19196019 | D431 | 2021-11-23 | B.1.160 | 98.70 % | Illumina, NovaSeq |
| EPI_ISL_10816733 | D432 | 2021-11-24 | B.1.160 | 98.80 % | Illumina, NovaSeq |
| EPI_ISL_19196020 | D434 | 2021-11-26 | B.1.160 | 98.00 % | Illumina, NovaSeq |
| EPI_ISL_19196021 | D440 | 2021-12-02 | B.1.160 | 93.20 % | Illumina, NovaSeq |

**Figure 2:**
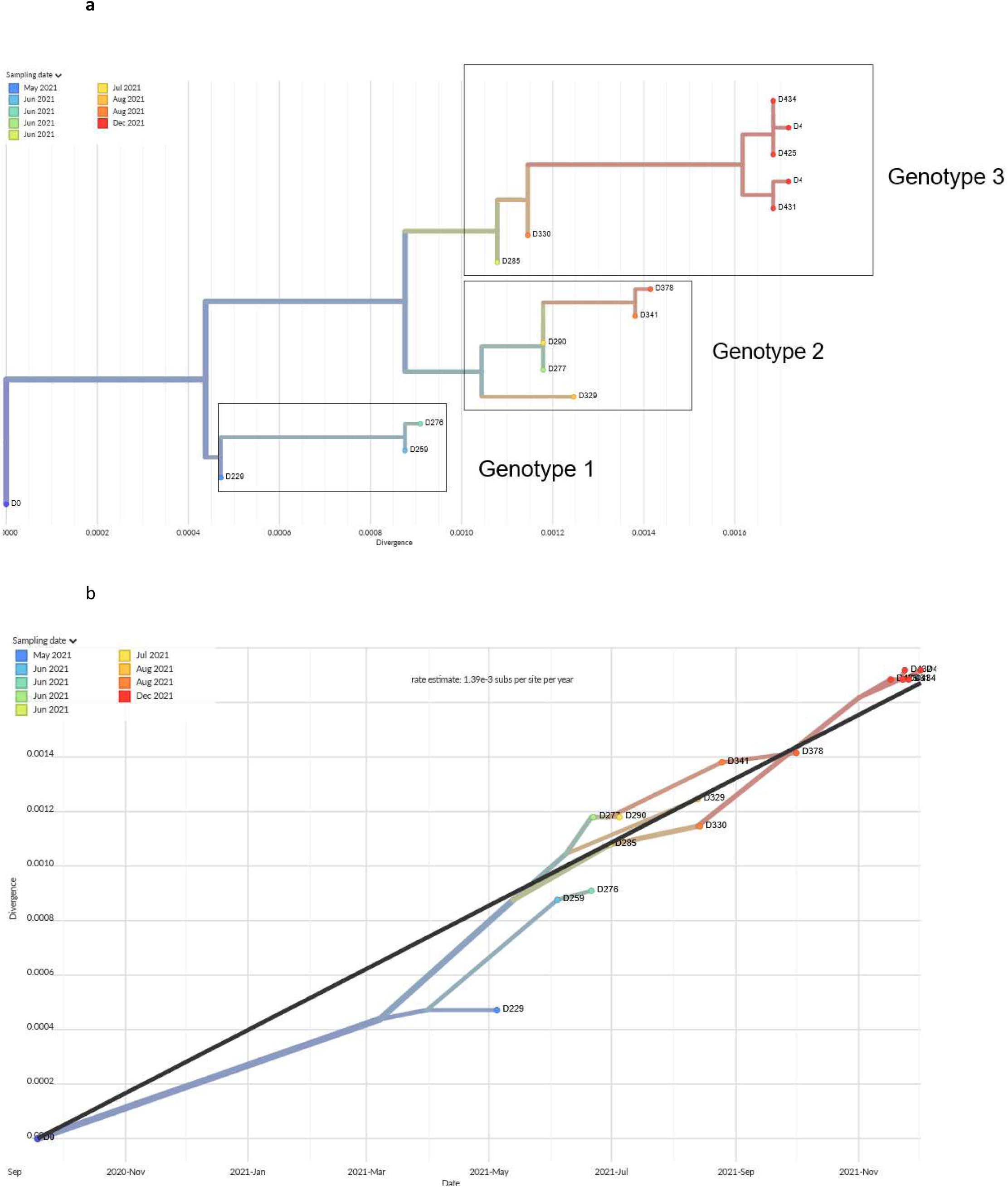
Within host evolution at consensus level. a) Maximum likelihood phylogenetic tree constructed with Nextstrain, based on the 16 whole SARS-CoV-2 genomes obtained from the immunocompromised patient between September 2020 and December 2021. Branch lengths represent the extent of genetic divergence. The phylogenetic tree reveals the placement of the sequences into three distinct genotypic groups, highlighted by dotted lines. b) Maximum likelihood phylogenetic tree constructed using Nextstrain, based on the 16 SARS-CoV-2 genome sequences taken from the patient. The y-axis reflects genetic divergence, the tree shows the rate of evolution of the sequences over time.

The temporal overlap between these genotypes can be explained by the coexistence of viral subpopulations within the host, potentially replicating in different anatomical compartments, as observed in long-term SARS-CoV-2 infections^49,6,50^. At coverage 10x, the intra-host evolutionary rate was estimated at 1.39 × 10□³ substitutions per site per year, 1.54 times higher than the estimated inter-host rate of 9.02 × 10□□ substitutions per site per year (Figure 2b, S1). This value was derived by comparing our dataset with 700 randomly selected SARS-CoV-2 genomes from locally circulating variants (September 2020–December 2021) deposited in the GISAID database (https://gisaid.org/) (Figure S1). These findings align with previous reports demonstrating accelerated viral evolution in chronic infections^5,16,51^, especially as in this particular case, the evolutionary dynamic of SARS-CoV-2 was driven by recombinant parental mutations associated with the emergence of *de novo* mutations.

### Expanding host diversity analysis with iSNVs and SNPs

We determined the optimal sequencing depth for analyzing intra-host SARS-CoV-2 diversity by comparing genome-wide and gene-level coverage at 50x and 200x (Figure S3a-b). While 200x coverage, representing 10 times the reciprocal of the MAF (5%), initially seemed preferable^52^, it resulted in a loss of SNP information (30 fewer substitutions in SNP_TOT, Figure S3c) and a drop of ≈11% in genome-wide coverage (80.82% ± 10.90 at 200x vs. 91.47% ± 7.24 at 50x, Figure S3a). 50x was selected as the optimal threshold, ensuring both SNP retention and reliable iSNV detection (≥5% MAF, ≥5 reads support) (see Material and Methods). Samples D259 and D440 were excluded due to less than 80% genome coverage (Figure S3c).

We sought to capture the full spectrum of viral evolution within the host, combining the contribution of exogenous mutations with assimilation by recombination and evolution by *de novo* emergence process. Hence, we first categorized mutations based on their frequency, to distinguish transitional mutations from fixed mutations, using a 50% allele frequency (AF) threshold: intra-host single nucleotide variants (iSNVs; AF <50%) and single nucleotide polymorphisms (SNPs; AF ≥50%). Additionally, mutations were grouped into two pools: “Within-host” (WH): mutations arising exclusively within the host, and “Total” (TOT): including all mutations, independently of their origin.

At day 229, a notable increase in iSNV richness was notable, reaching its maximum in the iSNV_TOT pool with 45 iSNV detected, representing the active phase of recombination, where the introduction of recombinant genomic segments temporarily increased intra-host diversity (Figure 3b). After reaching this peak, iSNV richness declined in both pools, showing fluctuations over time, which may be attributed to sampling bias or subpopulation circulation. From day 290, the iSNV_TOT and iSNV_WH curves are nearly indistinguishable, which means a minimal contribution of recombination-derived parental mutations to the iSNVs pools beyond this time point. The accumulation of SNP_TOT ranged from 20 substitutions at the beginning of the infection to a maximum of 56 SNPs at later stages. The number of SNPs originating from recombination remained stable throughout the infection, with eight lineage-defining B.1.1.7 substitutions acquired (C913T, C3267T, A23063T, C23271A, C23604A, C23709T, T24506G, and G24914C), while six lineage-defining B.1.160 substitutions were lost (C18877T, G22992A, G25563T, C25710T, C26735T, and T26876C) (Figure S2). The SNP_WH and SNP_TOT pools exhibited a simultaneous increase starting from day 330, although a decline in the number of SNPs (TOT and WH) was observed between days 329 and 330. This drop can be attributed to uncovered genomic positions as well as recombined segments, which involve both exogenous mutations from parental genomes and mutations that have emerged within the host. Notably, a decrease in iSNV richness is observed at days 329 and 378 compared to other time points. This drop could reflect variations of stochastic effects in viral population dynamics, emphasizing the need for careful interpretation of temporal fluctuations in intra-host diversity.

**Figure 3:**
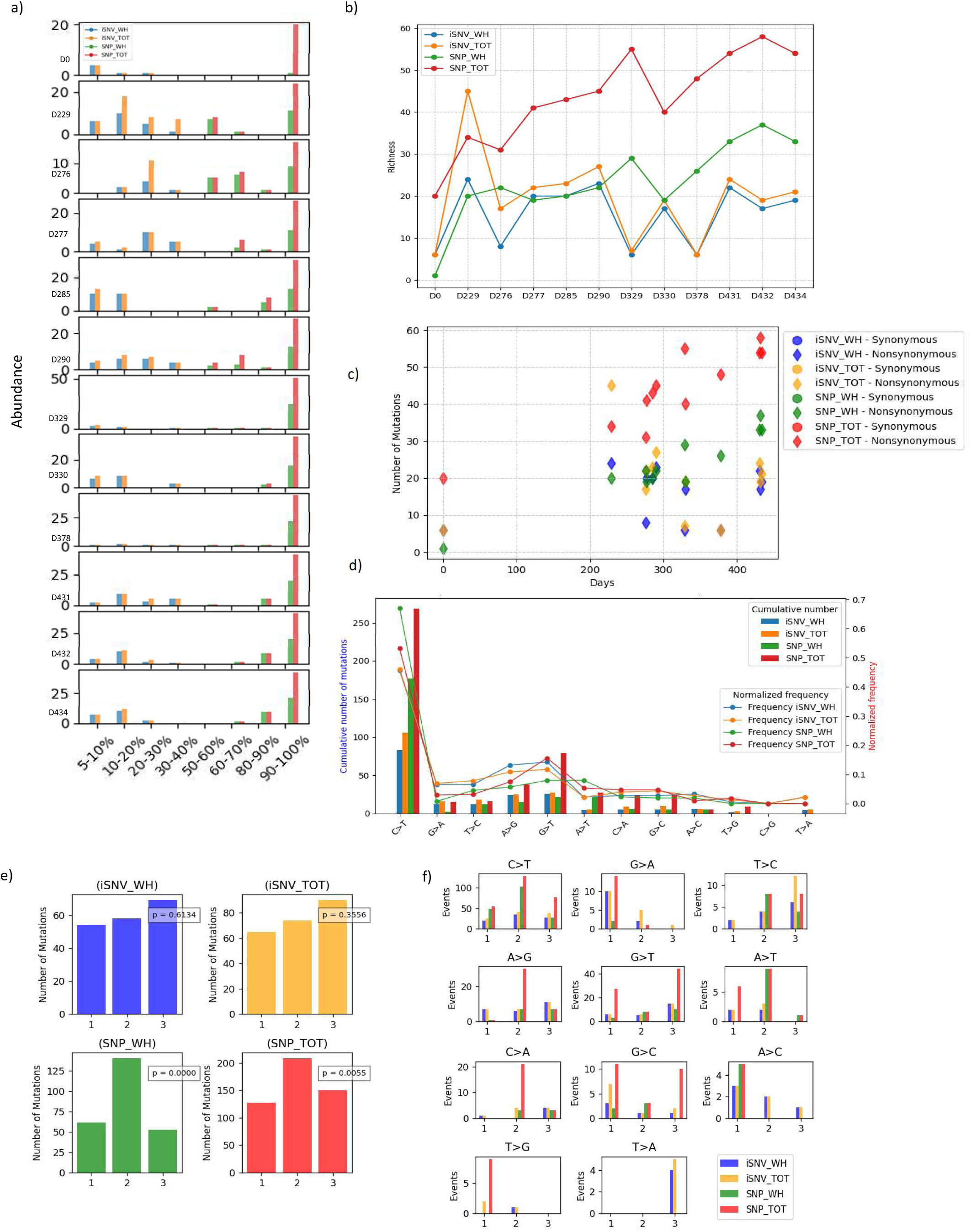
Distribution of iSNVs and SNPs: abundance, diversity, proportion of synonymous and non-synonymous mutations, and classification of mutation type. a)Abundance of iSNVs and SNPs illustrated by the distribution of mutations according to allelic frequency intervals (x-axis), membership of a mutation pool being indicated by a color code: blue for ‘within host’ iSNVs, yellow for ‘total’ iSNVs, green for ‘within host’ SNPs and red for ‘total’ SNPs, samples are indicated next to the y-axis. b) Richness represented by the number of mutations in the different pools as a function of time. c) Temporal accumulation of synonymous and non-synonymous mutations, classified as iSNV_WH (within host), iSNV_TOT (total), SNP_WH (within host) and SNP_TOT (SNP_TOT). d) Distribution of substitution classes in the four pools, displaying cumulative counts and normalised frequencies representing the proportion of each mutation class to the total number of mutations in each pool. e) Stacked bar charts showing the distribution of mutations at codon positions (1st, 2nd and 3rd) for each substitution type in the four pools. f) Number of mutations per codon position for each pool, with χ² test results and p-value.

The allele frequency distribution (Figure 3a) shows contrast in evenness between iSNVs and SNPs. SNPs (WH and TOT) predominantly clustered within the 90-100% frequency range, indicating low evenness, exceptions were observed at day 229 (D229), D276 and D290, where SNPs exhibit a broader distribution across the 50-100% frequency intervals. In contrast, iSNVs show a more dispersed frequency distribution, spanning a wider range of frequency bins than SNPs, particularly in samples D229, D277, D290, D330, and D431. The fluctuations in iSNV frequency distribution can be attributed to sampling bias or circulation of viral subpopulations. In contrast, the tighter clustering of SNPs (TOT and WH) within high-frequency bins reflects the stabilization and strong fixation over time.

We then examined the proportion of synonymous and non-synonymous mutations in the four pools (Figure 3c-S4). Non-synonymous mutations were significantly more frequent in the SNP pools (SNP_WH and SNP_TOT), while they were more frequent without reaching statistical significance in the iSNV pools (iSNV_WH and iSNV_TOT) (Figure S4). This more balanced distribution of non-synonymous and synonymous mutations may reflect the influence of genetic drift on transient mutations, where mutations arise stochastically and are not necessarily subject to immediate selective pressures.

Next, we analyzed the mutational spectrum of iSNVs in search of special signatures associated with pools. Across all pools, 11 different types of mutation were observed. A massive enrichment in C>T transitions was evident in all pools (Figure 3d), consistent with APOBEC-mediated RNA editing, a major contributor to SARS-CoV-2 evolution^53,36,37^. Additionally, the G>T transversion and A>G transition emerged as the second and third most frequently observed mutations in SNP_TOT and iSNVs, respectively. The high prevalence of G>T mutations in SNP_TOT is particularly noteworthy, as previous studies have highlighted its frequent occurrence and its association with oxidative damage-induced mutagenesis^54^, a process that can influence SARS-CoV-2 evolution within individual hosts. This is further supported by the second most prevalent G>T mutations in iSNVs, suggesting its active role in intra-host viral diversity. Furthermore, the A>G transition, which is also prominent in iSNV pools, has been linked to ADAR deaminase activity, another host-driven mechanism affecting viral RNA editing^55^. The mutation types T>C and G>A were slightly over-represented in iSNVs, and may result from ADAR and APOBEC activity respectively, acting on the complement strand on double-stranded RNA^56^.Given that SARS-CoV-2 RNA is mostly present in its positive-strand form, the C>T transition dominates, while G>A, which reflect APOBEC activity on the negative strand, is less prevalent^57^. Collectively, these three mutation types, C>T, G>T, and A>G, along with the more sporadic yet notable presence of T>C and G>A in iSNVs constitute the dominant mutational signature, underscoring the host antiviral machinery as a driving force in within host evolutionary landscape^58,59^.

We next examined the distribution of mutations across codon positions (Figure 3d-e). In iSNV pools, the distribution appeared relatively homogeneous, with no codon position standing out statistically, but a slight dominance of 3rd position, known for tolerating synonymous substitutions, with regard to the redundancy of the genetic code. In SNP pools, mutations preferentially affected the second codon position, leading to a significantly heterogeneous distribution (p < 0.05; Chi-square test). This preference was most pronounced in SNP_WH. The more balanced distribution observed in iSNVs is consistent with genetic drift, reflecting a more stochastic process where mutations accumulate as a reservoir of diversity without strong selective constraints. In contrast, the significant enrichment of mutations in the second codon position in SNPs suggests that selection is actively shaping mutation retention. This observation aligns with the fact that second-position mutations are more likely to cause amino acid changes, driving non-synonymous substitutions and potentially influencing viral adaptation.

When analyzing mutation type relative to codon position (Figure 3f), we similarly observed a more even distribution of mutations across positions for iSNV pools, particularly for C>T, A>G, G>T, G>C, and A>C mutations. In contrast, SNP pools exhibited a more localized distribution: for instance, G>A, T>G, and A>C mutations predominantly occurred at the first codon position, while C>T mutations were heavily biased toward the second position in SNPs. In iSNV pools, C>T mutations were more dispersed, occurring across all three codon positions. This visualization highlights fundamental differences in mutational preferences between transient within-host variants (iSNVs) and fixed mutations (SNPs). The preferential localization of SNPs in specific codon positions, particularly at the second position, strongly suggests that selection is actively shaping the fixation of certain mutations, while iSNVs act as a more neutral reservoir of genetic diversity.

Across all four mutation pools, a broad distribution of C>T mutations was observed across different nucleotide contexts (Figure 4,S4). In SNP_TOT and SNP_WH, C>T occurred in 16 and 17 different contexts, respectively, with the most represented being ACA, ACT, GCT in the two pools with TTC and CCT also more prevalent in the SNP_TOT pool. In iSNV_TOT and iSNV_WH, C>T is present in 21 and 20 contexts, primarily ACA, ACT, TTC. The selection process may shape the representation of C>T contexts in SNPs, potentially reflecting the mutational signature of APOBEC enzymes, which preferentially target single-stranded RNA in UC/AC motifs^57^.

**Figure 4:**
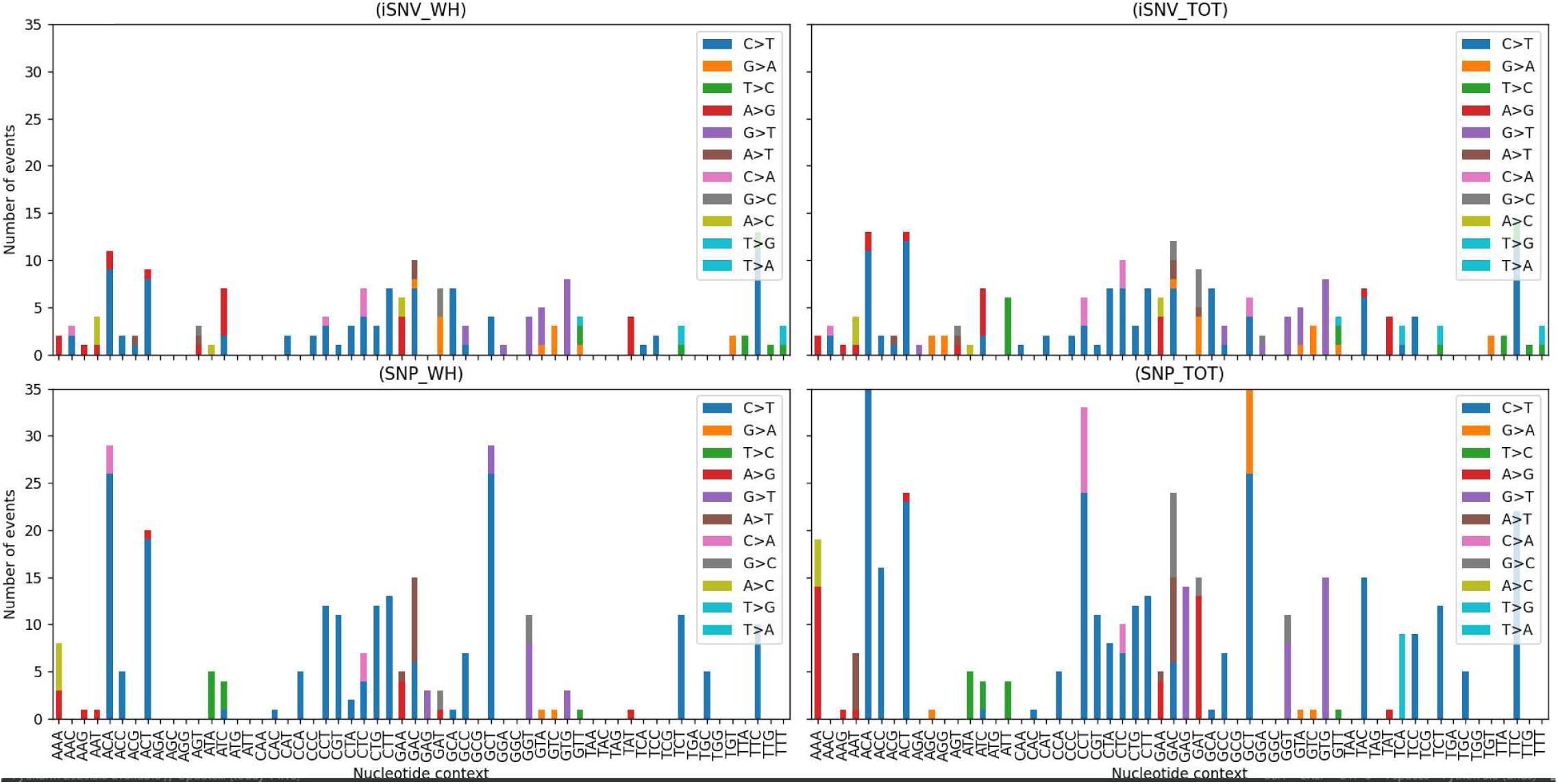
Trinucleotide context associated with mutation type across the four mutation pools: distribution and principal component analysis. Stacked bar plots showing the distribution of number of mutation types across different trinucleotide contexts within each of the four mutation pools: iSNV_WH, iSNV_TOT, SNP_WH, and SNP_TOT. Each bar represents the number of a given trinucleotide context, color-coded by mutation type.

In addition, A>G mutations, associated with ADAR activity, exhibit distinct context preferences. In iSNV_WH, the ATC, TAT, and GAA motifs were the most represented, consistent with ADAR’s tendency to target adenosines flanked by uridines or enriched in adenines, or eventually sensitive to GAA motifs^60^. In SNP_WH, A>G mutations predominantly occurred in AAA and GAA. This further supports the role of host-driven RNA editing mechanisms in shaping SARS-CoV-2 intra-host diversity.

### Quantitative diversity

As a precursor to the pN/pS analysis, we examined Shannon entropy along the genome, which quantifies mutational diversity. The whole genome entropy was adjusted according to gene coverage (normalized entropy), cumulative values by mutation group are shown in Figure 5c. Entropy was significantly higher in iSNVs from both WH and TOT pools, reinforcing the transient nature of intra-host mutations. Notably, no significant differences were observed between iSNV_WH and iSNV_TOT or between SNP_WH and SNP_TOT (Mann-Whitney U test), suggesting that within-host emergence of mutation dominates the mutational landscape, while the contribution of exogenous mutations inherited from parental genomes remains more limited at the whole-genome scale.

**Figure 5:**
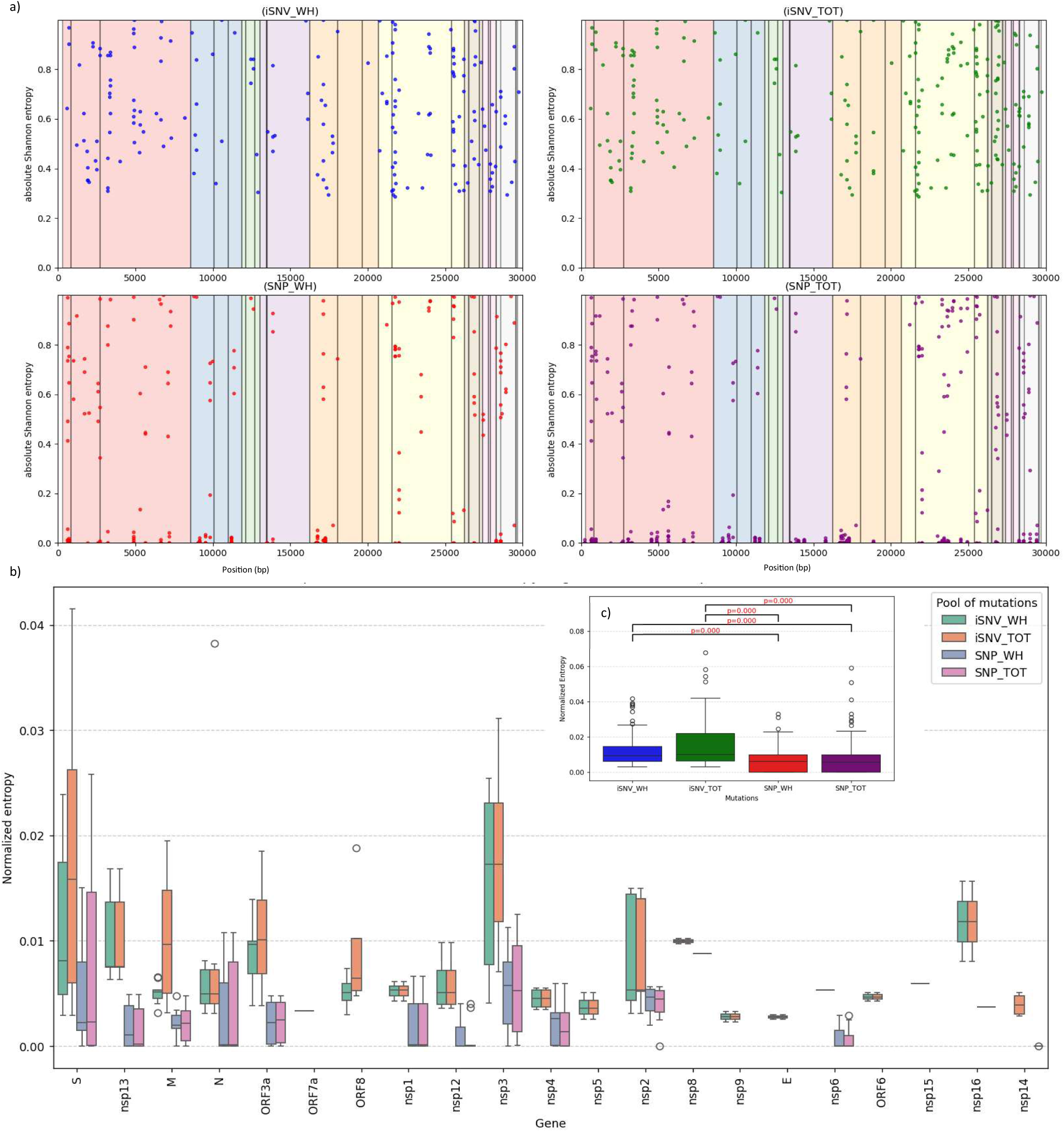
Gene level and wide genome normalized entropy. a)Scatter plots depicting the distribution of Shannon entropy along the viral genome with position (x-axis) within the four pools. The background colors highlight different genomic regions, including structural (S, N, E, M) and non-structural (ORFs, nsp) proteins. b)Boxplots show the normalized sequence entropy across different genomic regions for each mutation pool. c)Boxplots compare overall entropy distributions across the four mutation pools, with statistical significance denoted by p-values from Mann-Whitney U test.

We assessed further the impact of evolutionary dynamics at the gene level. Entropy was consistently higher in iSNVs than in SNPs (Figure 5a-b). This pattern reflects the greater transient diversity within the host before fixation.

Notably, the Spike (S) gene exhibited a distinct mutational pattern (Figure 5b). The lowest entropy was observed in iSNV_WH, suggesting strong purifying selection limiting intra-host diversification. In contrast, iSNV_TOT displayed the highest entropy, likely driven by recombination-mediated diversity. Additionally, SNP_WH exhibited low entropy with minimal variation, indicating that few iSNV_WH mutations reached fixation. In contrast, SNP_TOT showed higher entropy than SNP_WH, reinforcing the key role of recombination in shaping the Spike gene’s evolution. This suggests an initial phase of mutational influx in Spike (iSNV_TOT), followed by selection acting on beneficial variants (SNP_TOT).

Some genes exhibited low entropy across all pools, indicating strong fixation or evolutionary conservation. These included nsp5, nsp8, nsp9, E, nsp6, ORF6, nsp15, and nsp14.

The nucleocapsid (N), nsp13, nsp1 and nsp12 gene exhibited very low median entropy in SNPs; however, the observed variability (interquartile range) reflects the heterogeneity across samples (Figure S7). The high outlier in the iSNV-TOT pool for N and ORF8 corresponds to the active recombination phase, during which B.1.1.7 mutations were still present (at AF ≈15%) before being eliminated (Figure S7).

### Highly conserved genes

Highly conserved genes throughout the infection course were identified based on low pN/pS ratios, tolerance for high proportion of indeterminate pN/pS values (i.e., cases where both non-synonymous and synonymous mutations were absent, leading to an undefined ratio), and a very limited number of cases where pS = 0 and N > 3 (Non-synonymous), resulting in an infinite ratio (Figure 6 a-b,S6). Across all pools, several genes displayed low pN/pS ratios (M and nsp12) or no fixation of non-synonymous mutations (E, ORF6, ORF8, nsp16, nsp5, nsp9) with high undetermined ratio and no infinity value reported. This highlights a transient emergence of mutations, followed by their elimination, preserving the functional integrity of the proteins. NSP12, NSP7 and NSP8 form the RNA-dependent RNA polymerase (RdRp) holoenzyme^61^. These genes play critical roles in viral replication^61^. During this intra-host evolution, NSP12 retained its non-synonymous mutations inherited from the B.1.160 lineage, including G13993T, C14408T (ancestral), and G15766T (B.1.160). However, among the mutations that emerged within the host, only the synonymous mutation C13506T became fixed, while C13860T was only transiently fixed. This pattern suggests that the optimized changes in NSP12 from the B.1.160 lineage were preserved, while additional non-synonymous intra-host mutations did not persist, reinforcing the high conservation of NSP12 throughout this evolutionary trajectory, given its fundamental role in the SARS-CoV-2 life cycle.

**Figure 6:**
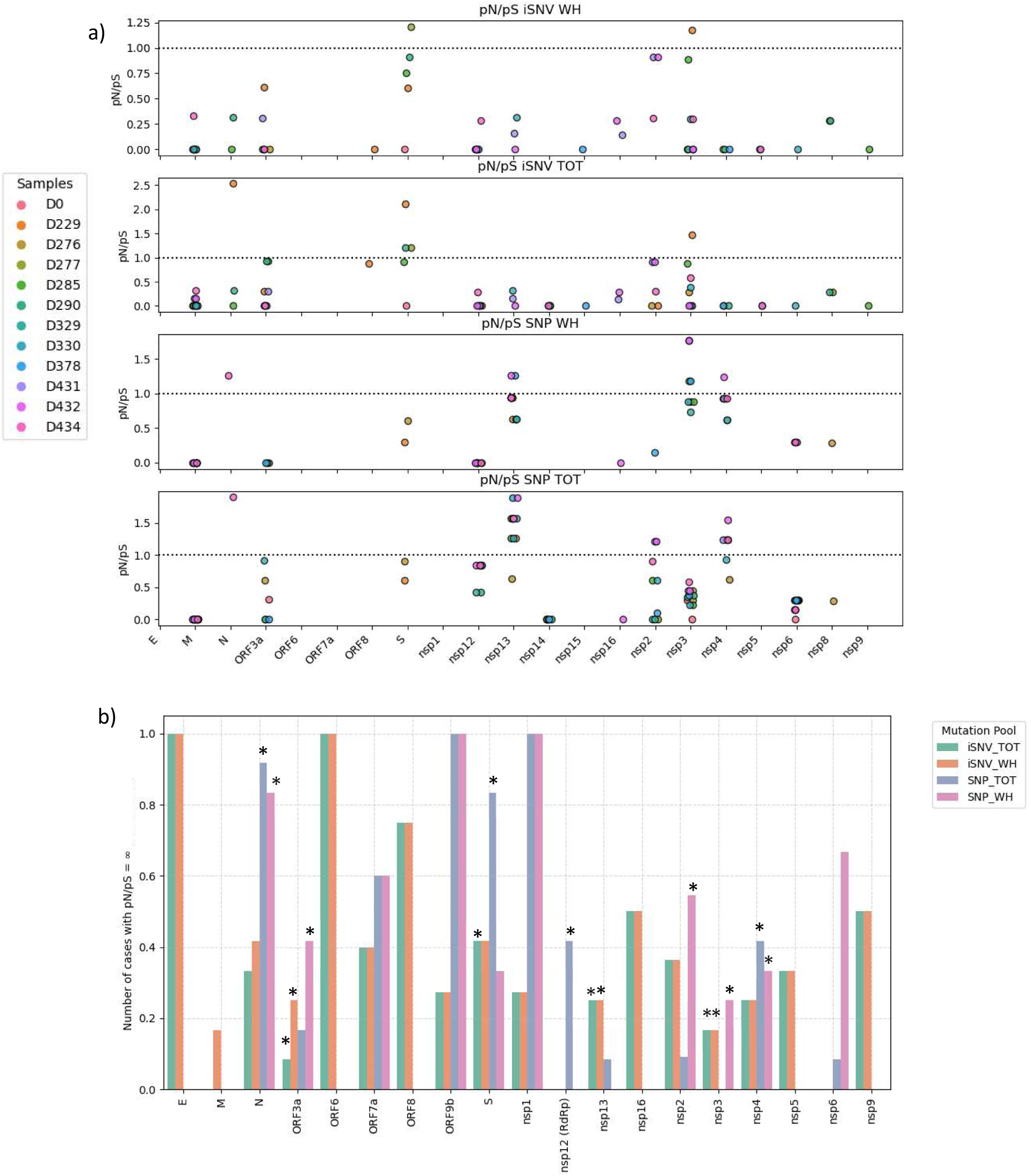
pN/pS ratio across SARS-CoV-2 genes in the four mutation pools. a)pN/S ratio calculated across different genomic regions of SARS-CoV-2, categorized into iSNV_WH, iSNV_TOT, SNP_WH, and SNP_TOT. Each panel corresponds to one of the four mutation pools b) Histogram representing the number of cases with pN/pS = ∞, referring to cases where pS=0 and N≥1, stars indicate pools where samples with pS=0 and N≥3 are found. c) Number of synonymous and non-synonymous mutations in Spike gene per sample according to the different pools.

Moreover, nsp14, nsp15, and nsp5 consistently displayed undetermined pN/pS values (pN = 0, pS = 0) in all pools, and sporadic low pN/pS values (Table 3, Figure 6a-S6). These genes are likely under functional constraints, where mutations that do emerge are rapidly eliminated by purifying selection. These arguments for nsp14 and nsp15 conservation reinforce their critical role in RNA proofreading and frequency of recombination^32^ respectively, which is relevant here.

**Table 3:** Evolutionary constraints and selection dynamics of SARS-CoV-2 genes across mutation pool.

| Gene | pN/pS | Infinite (N>3) | Indeterminate | Behavior across pools |
| --- | --- | --- | --- | --- |
| nsp1 | None | None | Present | Indeterminate |
| nsp2 | Variable: <1 in iSNV pools and SNP_WH, >1 in SNP_TOT in later samples | 3 samples in SNP_WH | Present (iSNV_WH and SNP_WH) | Few synonymous mutations in iSNV_WH/SNP_WH, positive selection in SNP_TOT at the final stage (global evolution process) |
| nsp3 | Variable: pN/pS<1 in SNP_TOT, some samples >1 in SNP_WH and iSNVs | Present in iSNVs and SNP_WH | None in SNP_TOT | Strong evolutionary constraint in SNP_TOT, sporadic positive selection in iSNVs/SNP_WH, occasional infinite values |
| nsp4 | Variable: low in iSNVs, >1 in some SNP_WH and SNP_TOT samples | Only in SNPs | Present in iSNVs | Positive selection in SNPs, accumulation of beneficial non-synonymous mutations |
| nsp5 | Very low | None (<3 non-synonymous mutation) | High | Conserved protein, possible purifying selection |
| nsp6 | Very low (pN/pS<1 in 6 SNP_TOT samples, sporadic <1 ratio in SNP_WH/iSNVs) | None (<3 non-synonymous mutation) | High in iSNVs | Highly conserved, purifying selection in SNPs and iSNVs (linked to autophagy interactions) |
| nsp7 | None | None | None | Indeterminate |
| nsp8 | Low, sporadic in iSNVs and SNPs | None | High | Indeterminate, signs of purifying selection |
| nsp9 | Low in iSNVs | None (<3 non-synonymous mutation) | High | Indeterminate, signs of purifying selection |
| nsp10 | None | None | None | None |
| nsp11 | Not covered | Not covered | Not covered | Not analyzable |
| nsp12 | Very low in all pools | Sporadic only in SNP_TOT | None in SNP_TOT, few in other pools | Highly conserved, negative selection since iSNV stage |
| nsp13 | Low in iSNVs, moderate in SNP_WH, high in SNP_TOT (>1 in 8 samples) | Sporadic in iSNVs only | None in SNP_TOT, few in other pools | Positive selection in SNPs (accumulation of non-synonymous mutations), negative selection in iSNVs |
| nsp14 | Very low in SNP_TOT | None | High in all pools (moderate in SNP_TOT) | Indeterminate, signs of purifying selection |
| nsp15 | Very low, sporadic in iSNVs (1 sample with ratio <1) | None | High | Indeterminate, signs of purifying selection |
| nsp16 | Very low, sporadic in iSNVs (2 samples <1 in iSNVs, 1 sample <1 in SNPs) | None (<3 non-synonymous mutation) | High | Indeterminate, signs of purifying selection |
| S (Spike) | Variable: 3 samples with ratio >1 in iSNV_TOT, 4 samples with ratio <1 in iSNVs_WH, 2 samples | Strongly present in SNP_TOT | Few in SNP_WH | Signs of positive selection in SNP_TOT: Impact of recombination-driven diversity. Minor contribution |
| M | Very low in all pools | None (<3 non-synonymous mutation) | Few | Highly conserved, purifying selection in SNPs and iSNVs |
| E | None | None (<3 non-synonymous mutation) | High | Indeterminate, signs of conserved protein |
| ORF3a | Very low in all pools | Few in iSNV_WH, iSNV_TOT and SNP_WH | Few | Highly conserved in SNP_TOT, but signs of positive evolution within the host. |
| ORF6 | None | None | High | Indeterminate, seems preserved |
| ORF7a | Not covered | Not covered | Not covered | Not analyzable |
| ORF8 | Low, sporadic in iSNV_WH and iSNV_TOT (1 sample with ratio <1) | None (<3 non-synonymous mutation) | Few | Indeterminate, signs of conserved protein |
| ORF9b | None | High in SNPs | Few | Signs of positive selection in SNPs |
| N | Variable : sporadic ratio >1 for 1 sample in SNPs and iSNV_TOT, 2 samples with ratio <1 iniSNVs | High in SNPs | Moderate in iSNVs | Signs of positive selection in SNPs |

Similarly, nsp6, for which an indeterminate pN/pS ratio has been found on numerous samples in iSNVs (Figure S6), exhibited strong negative pN/pS ratio in SNPs. Consistent with the literature, in SARS-CoV-2 and other coronaviruses, nsp6 has been shown to be highly conserved both in terms of nucleotides and protein sequence^62^.

To notice, the coverage of some genes was poor, including ORF7a, nsp11 and ORF8 in particular, for which we were unable to obtain valuable information on their selection.

### Positive selection and adaptive evolution

We characterized genes under positive selection by a pN/pS ratio >1 and/or the presence of infinite pN/pS values associated with NS>3, to account for the accumulation of non-synonymous likely providing adaptive advantage.

While most SARS-CoV-2 genes were constrained by purifying selection, several exhibited a positive selection, particularly in SNP_WH and SNP_TOT pools, as expected.

One of the strongest signals of positive selection was observed in SNPs from nsp13 (helicase), a gene critical for viral replication and immune evasion. The nsp13 gene displayed variable selective pressures in iSNVs, with most samples exhibiting a pN/pS ratio <1, indicative of purifying selection, and two samples showing an infinite pN/pS ratio (S=0 and N>3) (Table S2, Figure S7). As the infection progressed, positively selected within-host mutations progressively reached fixation in SNP_WH, driving a pN/pS>1 (Figure 6b). This transition from negative selection and neutral drift in iSNVs stages to strong positive selection underscores nsp13 as a key hotspot for adaptive mutations in immunocompromised patients^63,64^.

Similarly, nsp4 (replication complex) and ORF9b (an interferon antagonist) displayed strong positive selection, indicating their possible involvement in immune evasion and viral replication optimization. ORF9b, in particular, exhibited infinite pN/pS values in SNP pools, with three non-synonymous mutations detected at D434. Notably, the mutations that appeared in ORF9b were of within-host origin and not of the parental genomes.

While initially under negative selection in iSNVs, the Nucleocapsid gene underwent strong positive selection in both SNP_WH and SNP_TOT pools, particularly in the late stages of infection (Figure 6a-b). This was evidenced by a high number of infinite pN/pS (N>3) values.

The accumulation of 8 non-synonymous mutations in iSNV_TOT visible on day 229 came from parental genome and recombination process eliminated these B.1.1.7 mutations. These results are consistent with the fact that N is considered a vital hotspot for mutations^65^.

Initially, nsp2 showed negative selection or had many samples with undetermined values in iSNVs. Over time, it shifted to positive selection in SNP_WH, with three samples carrying four or more non-synonymous mutations and no synonymous mutations. Similarly, three SNP_TOT samples showed a pN/pS ratio >1, indicating positive selection emerging late in the infection (from day 431 onward), possibly linked to immune evasion.

At the iSNV_WH stage, only the non-synonymous mutation C2706T was detected, while all other mutations in SNP_WH appeared directly at >50% allele frequency (AF). This suggests a rapid evolutionary process, though it may also result from sampling limitations or transient mutations not being captured.

### The case of nsp3: selection analyzed on two levels

During its 440 days of evolution, nsp3 retained three synonymous mutations inherited from its parental genomes: S-C3037T (common ancestor), S-C4543T (B.1.160 lineage), and S-G5629T (B.1.160 lineage). As a result, purifying selection dominated in SNP_TOT (pN/pS <1), largely due to the fixation of these synonymous mutations. However, when focusing exclusively on SNP_WH and iSNVs, positive selection was detected (Table 3, Figure 6a-b). This suggests a fast evolutionary process, where non-synonymous mutations are arising and selected within the host, probably providing an adaptive advantage and consequently are reaching fixation. Hence, nsp3 exemplifies the case in which positive selection favors specific mutations within the host, increasing their allele frequency, while purifying selection prevails still after fixation due to the high number of inherited synonymous mutations.

### Optimized selection in the Spike protein

Spike exhibited strong positive selection in SNP_TOT pool, accumulating 6 to 8 non-synonymous mutations over the course of infection while no synonymous mutations were detected, leading to an infinite pN/pS ratio—except for the within host L821L, observed at D229 and D276 (Figure 6b). This suggests that positive selection favored amino acid changes leading to their persistence over time. The evolutionary trajectory of S indicates that mutations inherited from the parental genome B.1.1.7 were strongly selected and retained, while within-host mutations were less contributive. The iSNV_WH pool exhibited variable pN/pS, sporadically >1, and signs of positive selection (Figure 6b). This positive selection seemed weak on promoting the transition from iSNV_WH to SNP_WH; less sign of positive selection was observed for SNP_WH, highlighting a strong purifying filter that eliminated non-optimal variations.

This pattern suggests that recombination-introduced mutations had already undergone prior filtering, leading to a highly optimized combination in SNP_TOT. Spike transitioned from genetic drift in iSNVs (negative selection with occasional positive selection in iSNV_WH) to strong positive selection in SNP_TOT, reinforcing the idea that only the most functionally advantageous mutations were fixed, shaping a highly refined adaptive profile.

### Gene-scale optimized Spike configuration

During this evolution time period, five within host SNPs have arisen in the Spike (L5F, D796H, L821L, T618K, T572I), and some were evolutive convergence events (Figure 7a). L5F, a potentially CTL escape mutation, is located at a site recognized by Human Leukocyte Antigen (HLA) molecules, which are key players in presenting viral peptides to T cells for immune recognition^66^. L5F has been found in numerous VOCs such as B.1.177 or B.1.526 and has been shown to increase viral infectivity^67^. This mutation arose in parallel with D796H in the S2 Spike subunit at the first stage of infection, before being replaced by the Spike B.1.1.7 gene during the recombination process. D796H, a signature mutation of the variant of concern B.1.3.118, appeared within the host in the first described immunocompromised long term shedding patient^5^, following convalescent plasma therapy. As the patient received conva-lescent plasma therapy before the detection of this mutation (Figure 1a), it is possible that we faced a similar situation, promoted by exogenous antibodies. After recombination of the original Spike gene, the mutations T618K and T572I arose gradually (Figure 7a). T572I is a convergent mutation also present in the AY.32 variant while T618K is a rare substitution. The consensus total mutations observed in the Spike protein are illustrated in Figure 7b.

**Figure 7:**
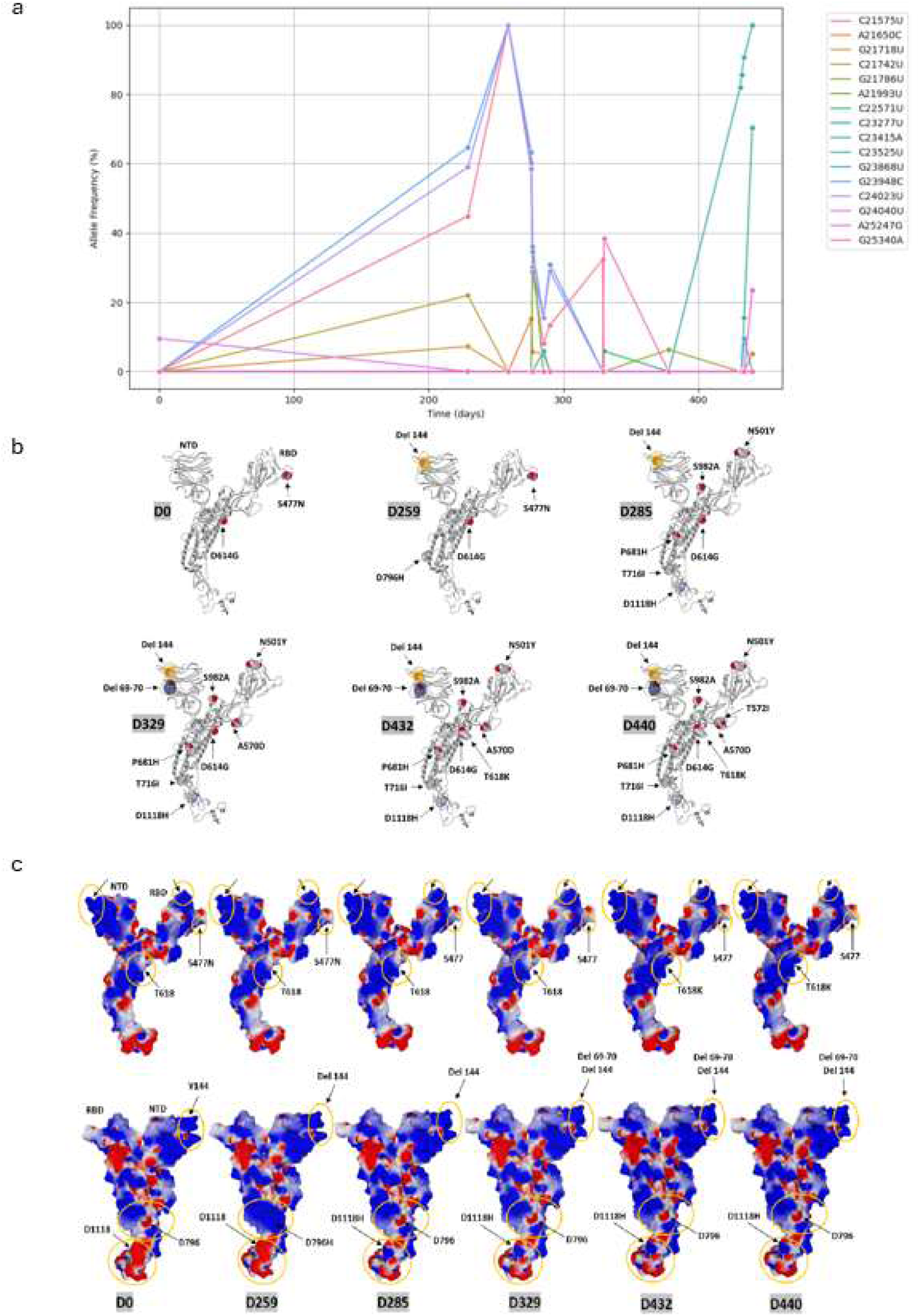
Evolution of allele frequencies of *Spike* within host substitutions, localization of total consensus mutations in the SARS-CoV-2 Spike protein and electrostatic potential of SARS-CoV-2 Spike proteins. a)Allele frequencies (%) of various within host Spike substitutions in the viral population across different time points (days). Each colored line represents a different mutation, with transient emerging SNPs (L5F, D796H, L821L, T618K, T572I) labeled. These mutations show distinct temporal patterns of allele frequency, reflecting their rise and fall in the viral quasispecies population over the course of the infection. b) The molecular structure of each mutant protein (from Day 0 to Day 440) is represented in grey ribbons and the mutations in atomic spheres (blue, nitrogen; grey, carbon; white, hydrogen; red, oxygen). Deletions are highlighted in orange (Del 144) and blue (Del 69-70) disks, taking as reference the last amino acid before the deletion (respectively V143 and I68). The signal sequence (amino acid residues 1-13) does not appear in the structure. NTD, N-terminal domain; RBD, receptor binding domain. c) The electrostatic potential of each mutant protein (from Day 0 to Day 440) is represented on the molecular surface (blue, positive; red, negative; white, neutral) in two opposite views. The upper panel corresponds to the same orientation of the Spike proteins as in Figure JF1. The lower panel shows the rear view obtained after a 180° rotation. Several regions are highlighted by orange circles to better visualize the changes associated with mutations.

### Molecular-scale optimized Spike configuration

As part of our analysis, we also studied the surface potential to gain a clear understanding of the protein-level impact and cell interaction resulting from the recombination and within host mutations observed (Figure 7c). Regarding within-host SNPs, an increase of electropositive surface is associated with T618K (upper panel) and D796H (lower panel), no additional significant changes have been observed regarding within host mutations. A first recombination event is visible at D259, in the N-terminal domain (NTD), where deletions 144 and 69-70 (from lineage B.1.1.7) appear, decreasing the NTD surface area facing the host cell plasma membrane (Figure 7c lower panel). This optimizes the virus’s initial interaction with lipid rafts^68^. The NTD condensation induced by deletions 144 and 69-70 is clearly visible in the lower panel of figure 6. At D285, the B.1.1.7 Spike variant is fully recombined, and the N501Y mutation is observed in the receptor-binding domain (RBD), permanently replacing the S477N mutation (present in B.1.160), thus increasing affinity for ACE2^69^. The rearrangement of the RBD surface is visible in the upper panel of figure 7. Other mutations include D614G, common to both variants B.1.160 and B.1.1.7, which facilitates the conformational unmasking of the RBD in the trimeric Spike^70^, S982A, which stabilizes this unmasked “up” conformation of the RBD, A570D and P681H, which may alter the Furin cleavage site. The D1118H mutation has a stabilizing effect on SARS-CoV-2 virions^71^. The NTD condensation induced by deletions 144 and 69-70 is clearly visible in the lower panel. The RBD surface rearrangement induced by the N501Y mutation is visible in the upper panel. Recombination has thus fixed a new assembly of upstream-optimized mutations, leading to subtle adjustments to the electrostatic surface potential. The deletions at positions 144 and 69-70 reduce the surface area of the NTD facing the host plasma membrane, optimizing the virus initial interaction with lipid rafts^47^. The absence of mutations linked to immune escape (e.g., K147 or R246) suggests that this patient did not produce neutralizing antibodies targeting the NTD, consistent with the patient’s immunocompromised status^72^ with IgG serology was negative as of May 25, 2021 (Table 1). The evolutionary significance of mutations in the Spike protein is closely related to their structural localization. While the NTD reshaping enhances its electrostatic binding to negatively charged membrane components, the RBD mutations (S477N and N501Y) increase Spike’s affinity for the ACE2 receptor, with N501Y providing a stronger advantage over S477N^69^). This recombination-driven selection likely optimizes virus-cell interactions, independent of immune escape processes, demonstrating the evolutionary benefits of such mutations.

### Convergent detection

Given the potential for SARS-CoV-2 long-term infection to generate new variants^51^, we investigated if within host SNPs selected for having reached or exceeded a 70% allele frequency, indicating for a strong tendency toward fixation, could be found in the general population and could therefore represent evolutionary convergences. We searched the GISAID database for their prevalence worldwide (Figure 8, Table S3), while it is also important to bear in mind that these results are subject to limitations due to sampling bias and inconsistent submission of data to GISAID. 20.31% of the selected within host SNPs were found in VOC (Variant of Concern) signature mutations, and 2 substitutions, C26858T, a synonymous substitution present in the Membrane gene and A28271T, a non-coding region substitution, were found re-spectively 22% and 52% of all sequences. The massive proportion of the C26858T synonymous substitution could be the result of optimized codon usage. Although UUC is more optimized in terms of codon usage, with codon frequencies of 20.3 for UUC and 17.6 for UUU^73^, the widespread replacement by UUU may confer advantages related to RNA structure or stability, suggesting an adaptation during within-host viral evolution. This change might enhance viral fitness through mechanisms beyond simple codon optimization, such as improving RNA folding or influencing the regulation of translation. The mutation A28271T is in the N gene Kozak translation initiation box^74^ and the A to T substitution is responsible for a reduced protein expression of the ORF9b gene, and therefore increases the production of the N protein relative to the ORF9b protein. A similar effect was observed with the B.1.1.7 deletion A28270-^39^. The parental strain B.1.1.7 that infected the patient following the initial infection carried the A28270-deletion, within the Kozak sequence. However, this deletion was not maintained during the intra host evolutionary process, as it was detected at an allele frequency of 15% on day 229 (Figure S2) before being completely eliminated. The substitution A28271T, emerged at the same position, potentially restoring the functional advantage previously provided by the deletion.

**Figure 8:**
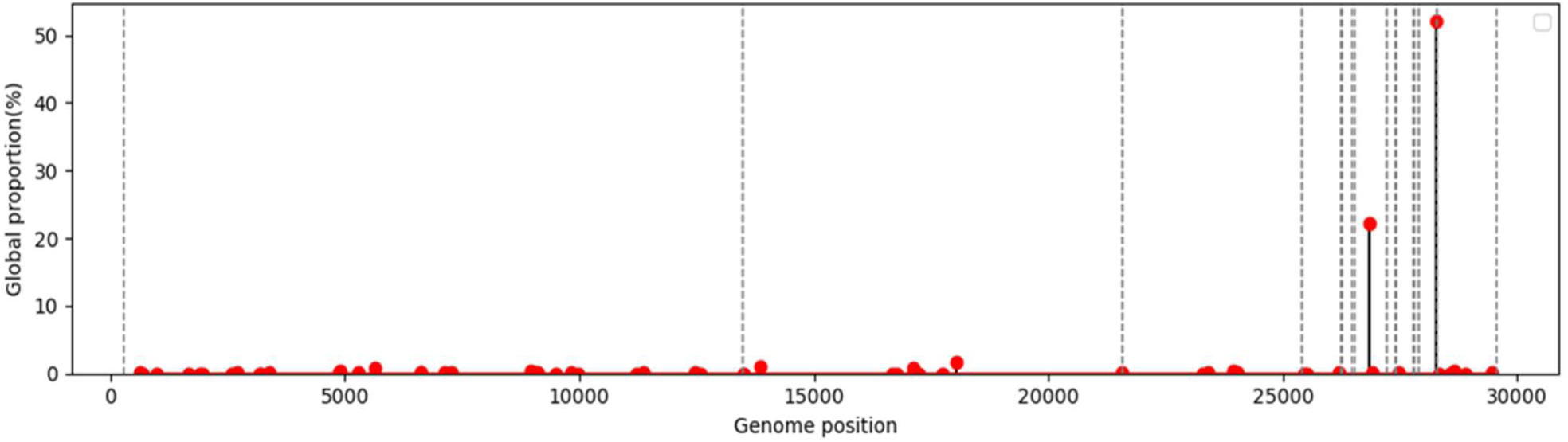
Global proportion of high fixed within host SNPs based on GISAID sequences. Graph showing the proportion (%) of within host SNPs, selected for having reached or exceeded a 70% allele frequency, observed in the patient among the general population, at different positions in the viral genome.

### The wild type-like restoration of ORF3a and M

From D0 to D329, the two parental mutations remained detectable but were subsequently eliminated (Figure S7). While this could result from a reversion event, it is more likely due to recombination, given that five mutations disappeared completely and did not reappear in later samples: NS-25460T (WH), S-G25563T (B.1.160), S-C25710T (B.1.160), NS-C26204T (WH). Following this elimination, only two additional mutations—one synonymous (S) and one non-synonymous (NS), both within-host (WH) transiently reached the fixation threshold (50%) before disappearing: NS-C25517T, which peaked at 61.9% (D432) before being lost and S-C25515A, which briefly reached 83.2% (D330) before vanishing. This case represents a scenario where recombination facilitated a reversion to the WT ORF3a.

For the Membrane gene, we observed a similar situation: the parental B.1.160 S-C26735T and S-T26876C mutations were eventually eliminated at day 434 and the S-C26858T (WH) reached fixation.

### Impact of intra-host evolution on the ORF8 gene

The accessory protein ORF8 plays a crucial role in the pathogenesis of SARS-CoV-2 due to its secretory nature and high plasticity, which allow it to interact with a wide array of host proteins, making it a key player in both viral replication and immune evasion^75,76,77^. In sample D378, a PCR analysis (Figure 9a) of the genomic region spanning 24,813-29,074 revealed a 126-nucleotides deletion from positions 27,915 to 28,040 (see supplementary Material and Methods). This within host deletion, which gradually became the dominant variant after 378 days of infection resulted in a loss of 42 amino acids, accounting for 34.43% of the ORF8 protein. Similar truncated forms of ORF8 have been observed in variants such as B.1.1.7, where a stop codon was introduced^76^. This 126-nt deletion in ORF8 was fortuitously discovered following a targeted PCR covering this region (Figure 9a) but was initially identified as an uncovered region in consensus genomes (Figure 9b). The deletion was obtained by the read assembly (Figure 9b), which was able to circumvent the analysis bias caused by the use of 50 bp read lengths in the mapping strategy. Indeed, in mapping approaches, reads are directly aligned to a reference genome, serving as a backbone, and deletions are inferred from gaps or reduced coverage. If a deletion is larger than the length of the reads (for example, a 126-nt deletion compared with 50 bp reads), the reads covering the newly deleted region may not align correctly and may be rejected as ambiguous, leading to an uncovered region in the consensus genome. In contrast, *de novo* assembly reconstructs the genome without relying on a reference sequence, enabling the detection of novel structural variations such as large deletions. This issue mirrors the challenges identified by Brandt et al.^78^ who showed that a large deletion (168 nt) in ORF8 was not detected by variant calling pipelines and standard sequencing technologies.

**Figure 9:**
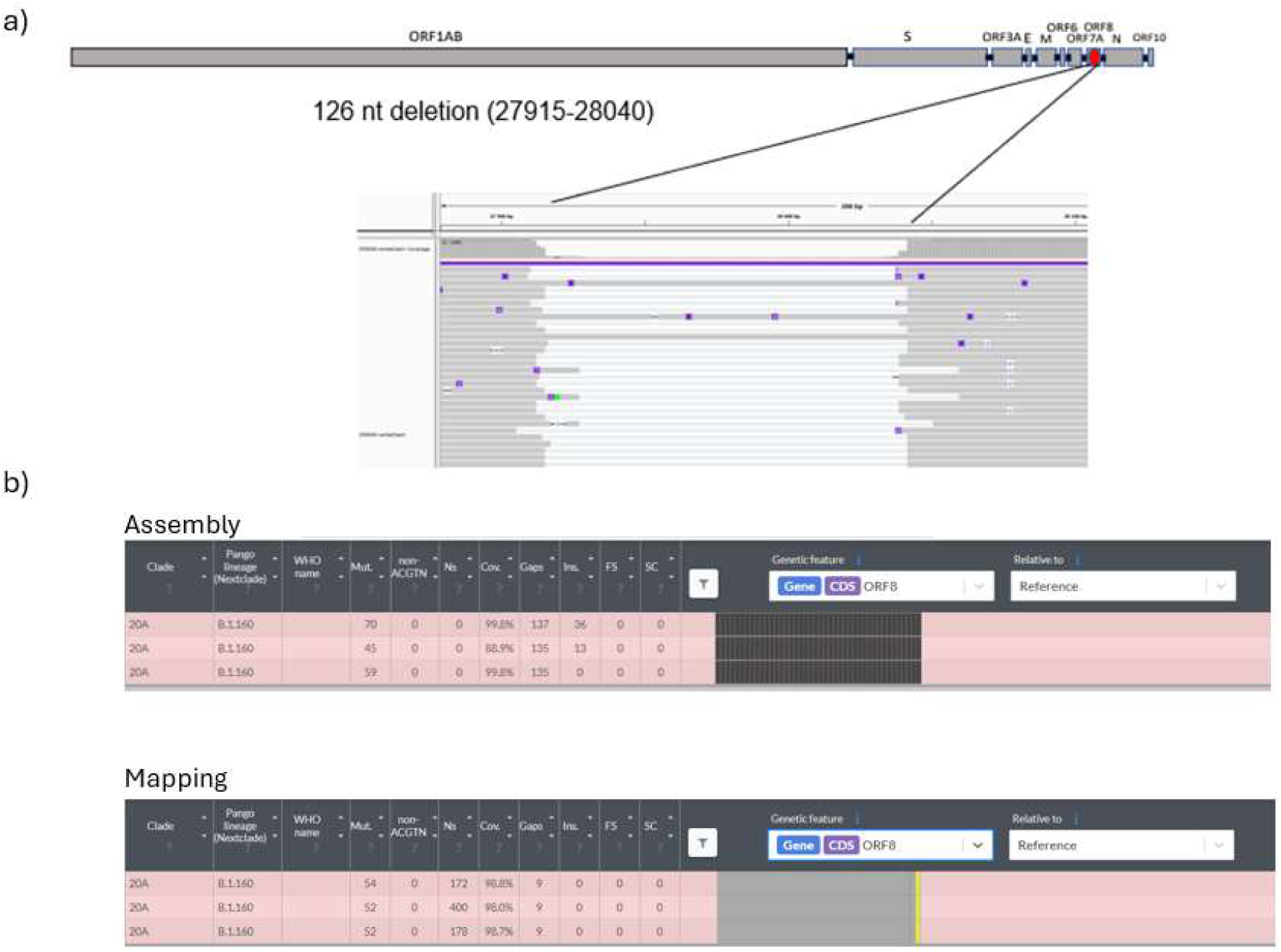
Emergence of ORF8 126-nt deletion since D378. a)Visualization of a PCR product sequencing on the region made on the sample D378. Mapped and sorted reads are visualized by IGV, a large nucleotide deletion is visible. b) Visualisation of the ORF-8 deletion on the consensus genomes obtained by assembly (top panel) and visualisation of the deletion detected as a missing region, obtained by mapping (bottom panel).

## DISCUSSION

This case study illustrates the dynamic evolution of SARS-CoV-2 within an immunocompromised patient, initially infected with the B.1.160 variant and later with B.1.1.7. The combination of immunodeficiency and the treatments received, such as convalescent plasma, created a favorable environment for accelerated viral evolution. The intra-host evolution rate observed in this case (1.39 × 10^-3 substitutions per site per year) was significantly higher than the inter-host rate (9.02 × 10^-4 substitutions per site per year), aligning with previous findings^5,16,51,79^ and relies only partially on recombination from parental genomes. The temporal overlap and coexistence of distinct genotypes within the host suggest the circulation of viral subpopulations across different anatomical sites^80,49,6,50^. Although this study is based on a single case, its significance is reinforced by the exceptionally long duration of the reported infection (440 days), making it one of the longest documented cases of SARS-CoV-2 persistence in an immunocompromised patient. The robustness of the analysis is further supported by the availability of high-quality samples (12 whole genomes sequences with more than 80% coverage at 50x depth).

However, our study has several limitations, the first being that as a retrospective study, we were not able to plan sequencing data points or analyzing other anatomical sites of interest such the lower respiratory system, as sample collection was dependent of previously banked specimens. Consequently, some key aspects of viral evolution, particularly the very initial stages of recombination could not be detailed. Reconstructing multiple recombination events through haplotype analysis, including within-host mutations, may have provided valuable insights and could represent an interesting working perspective, giving the challenge due to the short read length (50 bp), which restricts the ability to confidently link adjacent mutations within the same haplotype.

Another limitation concerns the restricted analysis of the immune response. Only three serological data points were available, which limited our ability to assess how immune pressure may have shaped viral evolution. A more comprehensive approach, including HLA typing and T-cell response characterization, could have provided deeper insights into the mechanisms driving the evolutionary processes at play. Since serology was negative from the end of May, evolutionary immune-pressure came essentially from innate and T-cell immunity since June. Once again, the retrospective nature of the study prevented us to explore these aspects in greater depth.

Patients with prolonged infections have been shown to accumulate more intra-host single nucleotide variants (iSNVs) than patients with shorter infections^63^, and should not be neglected as critical insights into the mechanisms driving intra-host viral evolution and the emergence of new variants^81,34,82,83^. Given the potential bias introduced by sequencing errors, we focused on iSNVs with allele frequencies between 5% and 50%, used iVar, a highly accurate tool for detecting intra-host single-nucleotide variants^84^ and applied optimal depth and quality thresholds to ensure reliable results.

In our study, some genes showed very few iSNV-WH but high SNP_WH, as they may have reached fixation too rapidly for detection, likely due to sampling limitations. Additionally, setting a MAF threshold at 5% may have restricted the ability to capture early-stage evolutionary dynamics, though this choice was necessary to ensure data reliability while minimizing false positives.

Stanevich et al^79^. demonstrated the emergence of mutations that facilitated immune evasion, including S:del141-144, associated with reduced antigen presentation by both HLA class I and II molecules, S:R273S, nsp3:T1456I and N:P6T, previously identified as immunogenic peptides targeted by T cells, which induced the loss of their ability to elicit T-cell responses. Comparing our findings with subsequent studies would help assess whether similar immune escape mechanisms could be at play in this prolonged infection.

The Envelope (E) and Membrane (M) remained highly conserved throughout the 440-day infection, suggesting strong functional constraints preserving their roles in viral assembly and exit. Similarly, nsp12 displayed high iSNV entropy but remained strongly constrained at the SNP level, aligning with its essential function in replication, where excessive mutations are often deleterious. Interestingly, E was previously reported to evolve within-host in another long-term infection case^79^ whereas in our study, it remained mostly conserved. This discrepancy could stem from selective pressures and HLA diversity in distinct host environments, warranting further investigation.

Several genes exhibited positive selection, including Spike, N (nucleocapsid), ORF9b, nsp13, nsp4, nsp2, and nsp3, highlighting their roles in immune evasion, and adaptation to the host environment.

N and ORF9b, known to interfere with type I and III interferon (IFN) responses, showed strong positive selection, reinforcing their role in modulating innate immunity^85,86^. Moreover, nsp2 underwent positive selection too, likely driven by interactions with IFN-β^87^ in the same way. Nsp13 (helicase) has been shown to accumulate mutation in immunocompromised patients^63^, possibly due to its role in host-pathogen interactions and immune modulation. Nsp3, which plays a role in innate immune modulation, displayed accumulated non-synonymous mutations, similar to findings by Stanevich et al.^79^, where mutations in this region led to altered recognition by T lymphocytes.

ORF3a is an abundantly accumulated accessory protein facilitating virus exit via lysosome deacidification^88,89^. Interestingly, recombination led to a reversion to a wild-type-like sequence, suggesting a selective advantage in retaining its ancestral functional state. The loss of multiple mutations in a coordinated manner suggests a genomic replacement event, likely driven by recombination, which may help restore a more genetically stable ORF3a^90^ configuration which needs further investigation giving that the ORF3a has been implicated in modulating the host immune response, notably by interacting with key proteins such as IKKβ and NEMO in the NF-κB signaling pathway^89^. This finding raises the possibility that some mutations associated with reduced pathogenicity, such as those seen in Omicron lineages^91^, could be reversible through recombination. Interestingly, the Membrane gene showed the same return to wild-type sequence by recombination. Additionally, since the E gene did not exhibit mutations from either parental strain, it is possible that it was not subject to recombination

The massively predominant C>T mutations encountered both, in iSNVs and SNPs (from within host and parental genomes origin) is known to be induced by the host APOBEC-mediated RNA editing family and has been extensively documented^92,54,93^in the global population and immunocompromised long carrier patients. APOBEC targets cytosine for deamination resulting in G>A mutations on the complementary strand during viral replication^94^. This mechanism, which involves the conversion of cytidines to uridines, likely contributes to increased viral plasticity, aiding the virus adaptation to the host^58,37^A>G and G>T were the two types the most represented among within host iSNVs. G>T mutations are frequently observed globally^95,96^ and are linked with oxidative stress caused by reactive oxygen species^59^. In addition, the A>G transition mediated by the ADAR protein has also contributed to the emergence of within host mutations, further emphasizing the importance of host-pathogen interactions in SARS-CoV-2’s intra-host evolution^55,97^. The dominance of A>G, C > T, and G > T mutations during intra-host evolution has been observed in cohorts across the world, suggesting that these molecular mechanisms are consistently at play in shaping SARS-CoV-2’s evolution ^81^.

Large deletions in ORF8 have also been reported in other lineages^98^. It is known that the presence of a full-length ORF8 is not essential for viral replication, and the loss of this gene may even confer a selective advantage by facilitating immune evasion or enhancing viral persistence. This phenomenon of “adaptive gene loss” has been described in SARS-CoV-2, where the loss of ORF8 can provide benefits to the virus, likely by reducing the immune system’s ability to detect and respond to infection^99^. This highlights the evolutionary flexibility of SARS-CoV-2, with ORF8 deletions serving as a potential adaptive mechanism during prolonged intra-host viral evolution.

Finally, this study emphasizes the need for comparative genomic analyses of SARS-CoV-2 in long-term carriers to deepen our understanding of the virus’s adaptive ability within the host. Gaining insights into host-virus interactions is crucial for elucidating the mechanisms that enable prolonged persistence, even in the absence of severe immunosuppression, as suggested in cases of long COVID^100,101,102^, a persistent global public health concern^101^. Understanding how the virus adapts at genomic, proteomic and structural levels will be instrumental in identifying novel therapeutic targets, ultimately contributing to the development of more effective treatment strategies as genomic diversity accumulation could act as a potential mechanism for viral persistence.

## Acknowledgments

We are grateful to the IHU NGS platform, Claudia Andrieu, Sofiane Bakhour, Amira Doudou, Idir Kacel and Vincent Bossi. We are also grateful to Jeremy Delerce for providing data on circulating variants of B.1.1.7. We gratefully acknowledge all the contributors of the sequences deposited in GISAID used on the global population SARS-CoV-2 tree.

## Author contributions

Conceptualization: EB, PP, PEF ; Contributed materials/analysis tools: EB, JF ; Analyzed the data: EB, JF ; Writing: EB, JF; review & editing: PP, PEF Supervision: PEF All authors have read and agreed to the published version of the manuscript.

