## Supplemental Material, Supplemental Tables and Figures 1-7 for "Long-Term Intra-Host Evolution of SARS-CoV-2 in an Immunocompromised Patient: Recombination and Within-Host Mutations Driving Viral Adaptation": supplementary.docx

SUPPLEMENTARY METHODS

PCR amplification of [24,813-29,074]

Briefly, RNA samples were extracted using the EZ1 Virus kit with an EZ1 Advanced XL instrument (Qiagen) following the manufacturer’s recommendations. PCR amplification of the [24,813-29,074] region was performed in a 25 µL total volume using the SuperScript III One-Step RT-PCR Kit (Invitrogen, Carlsbad, CA, USA), using the forward: ATGGAAAAGCACACTTTCCT; reverse: GCTTTAGTGGCAGTACGTTT primers at concentrations of 200 nM per reaction.

PCR was performed using the following conditions: 50 ◦C for 25 min, 95 ◦C for 2 min, then 40 cycles including 15 s at 95 ◦C, 45 s at 60 ◦C, and 2 min at 68 ◦C. Amplicons were sequenced using the Nanopore technology on a GridION instrument (Oxford Nanopore Technologies Ltd., Oxford, UK), following the manufacturer’s instructions. Raw reads were mapped on the SARS-CoV-2 reference genome using Minimap2 (https://github.com/lh3/minimap2), bam files were then filtered using SAMtools (v1.13; https://github.com/samtools/2008-2023 Genome Research Ltd.) combined with an in-house awk script.

TABLES

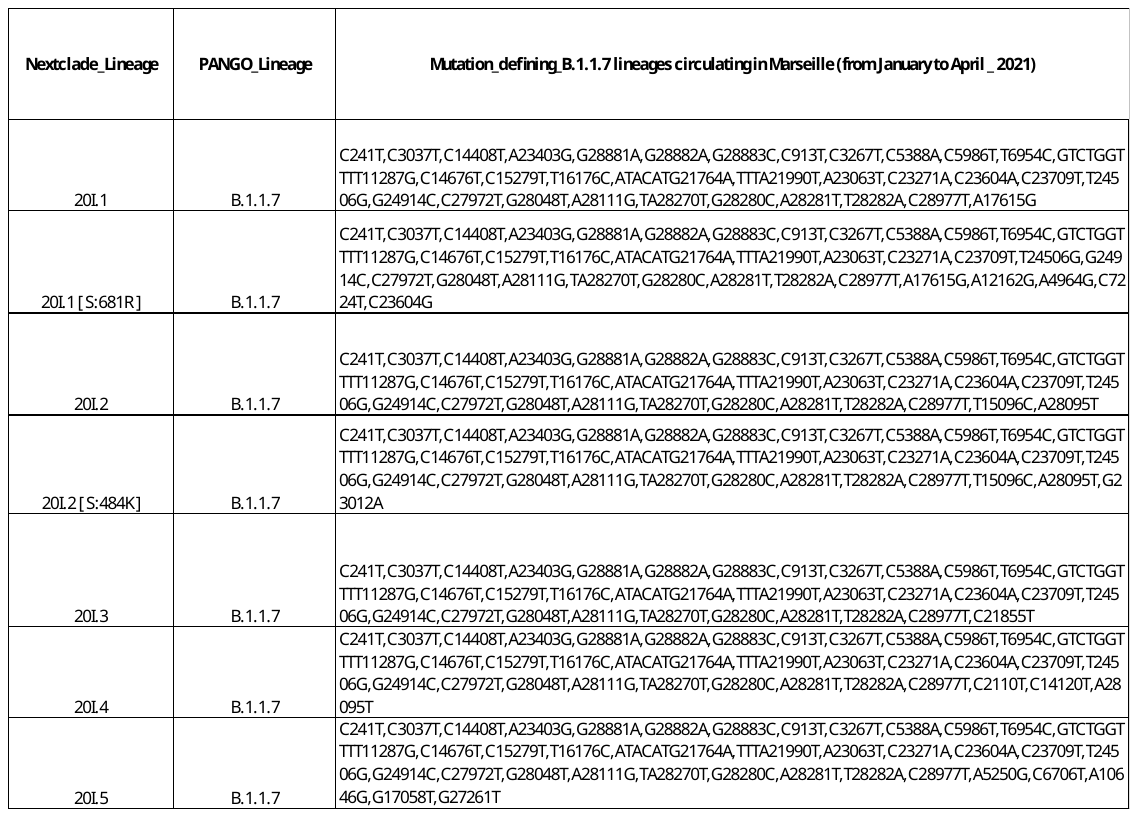

Table S1: List of mutations found in sequenced B.1.1.7 variants circulating in Marseille from January 2020 to May 2021

|  | **Sample** | | **Region** | **S_Mutations_iS**  **NV_WH** | | **NS_Mutations_ iSNV_WH** | | **S_Mutation s_iSNV_TOT** | | **NS_Mutations _iSNV_TOT** | | **S_Mutations _SNP_WH** | | **NS_Mutatio ns_SNP_WH** | | **S_Mutations _SNP_TOT** | | **NS_Mutations _SNP_TOT** |
| --- | --- | --- | --- | --- | --- | --- | --- | --- | --- | --- | --- | --- | --- | --- | --- | --- | --- | --- |
|  | **D0** | | **E** | 0 | | 0 | | 0 | | 0 | | 0 | | 0 | | 0 | | 0 |
|  | **D229** | | **E** | 0 | | 0 | | 0 | | 0 | | 0 | | 0 | | 0 | | 0 |
|  | **D276** | | **E** | 0 | | 0 | | 0 | | 0 | | 0 | | 0 | | 0 | | 0 |
|  | **D277** | | **E** | 0 | | 0 | | 0 | | 0 | | 0 | | 0 | | 0 | | 0 |
|  | **D285** | | **E** | 0 | | 0 | | 0 | | 0 | | 0 | | 0 | | 0 | | 0 |
|  | **D290** | | **E** | 0 | | 0 | | 0 | | 0 | | 0 | | 0 | | 0 | | 0 |
|  | **D329** | | **E** | 0 | | 0 | | 0 | | 0 | | 0 | | 0 | | 0 | | 0 |
|  | **D330** | | **E** | 0 | | 1 | | 0 | | 1 | | 0 | | 0 | | 0 | | 0 |
|  | **D378** | | **E** | 0 | | 0 | | 0 | | 0 | | 0 | | 0 | | 0 | | 0 |
|  | **D431** | | **E** | 0 | | 0 | | 0 | | 0 | | 0 | | 0 | | 0 | | 0 |
|  | **D432** | | **E** | 0 | | 0 | | 0 | | 0 | | 0 | | 0 | | 0 | | 0 |
|  | **D434** | | **E** | 0 | | 1 | | 0 | | 1 | | 0 | | 0 | | 0 | | 0 |
|  | **D0** | | **M** | 0 | | 0 | | 0 | | 0 | | 0 | | 0 | | 2 | | 0 |
|  | **D229** | | **M** | 1 | | 0 | | 3 | | 0 | | 1 | | 0 | | 1 | | 0 |
|  | **D276** | | **M** | 1 | | 0 | | 3 | | 0 | | 0 | | 0 | | 0 | | 0 |
|  | **D277** | | **M** | 0 | | 0 | | 2 | | 0 | | 1 | | 0 | | 1 | | 0 |
|  | **D285** | | **M** | 1 | | 0 | | 1 | | 0 | | 0 | | 0 | | 2 | | 0 |
|  | **D290** | | **M** | 1 | | 0 | | 3 | | 0 | | 1 | | 0 | | 1 | | 0 |
|  | **D329** | | **M** | 1 | | 0 | | 1 | | 0 | | 1 | | 0 | | 3 | | 0 |
|  | **D330** | | **M** | 1 | | 0 | | 1 | | 0 | | 0 | | 0 | | 2 | | 0 |
|  | **D378** | | **M** | 0 | | 0 | | 0 | | 0 | | 2 | | 0 | | 2 | | 0 |
|  | **D431** | | **M** | 0 | | 1 | | 2 | | 1 | | 1 | | 0 | | 1 | | 0 |
|  | **D432** | | **M** | 0 | | 1 | | 2 | | 1 | | 1 | | 0 | | 1 | | 0 |
|  | **D434** | | **M** | 1 | | 1 | | 1 | | 1 | | 1 | | 0 | | 1 | | 0 |
|  | **D0** | | **N** | 0 | | 0 | | 0 | | 0 | | 0 | | 0 | | 0 | | 2 |
|  | **D229** | | **N** | 0 | | 2 | | 1 | | 8 | | 0 | | 2 | | 0 | | 4 |
|  | **D276** | | **N** | 0 | | 0 | | 0 | | 0 | | 0 | | 2 | | 0 | | 3 |
|  | **D277** | | **N** | 0 | | 1 | | 0 | | 1 | | 0 | | 1 | | 0 | | 3 |
|  | **D285** | | **N** | 1 | | 0 | | 1 | | 0 | | 0 | | 1 | | 0 | | 3 |
|  | **D290** | | **N** | 1 | | 1 | | 1 | | 1 | | 0 | | 1 | | 0 | | 3 |
|  | **D329** | | **N** | 0 | | 0 | | 0 | | 0 | | 0 | | 2 | | 0 | | 4 |
|  | **D330** | | **N** | 0 | | 0 | | 0 | | 0 | | 0 | | 1 | | 0 | | 3 |
|  | **D378** | | **N** | 0 | | 0 | | 0 | | 0 | | 0 | | 1 | | 0 | | 3 |
|  | **D431** | | **N** | 0 | | 1 | | 0 | | 1 | | 0 | | 3 | | 0 | | 5 |
|  | **D432** | | **N** | 0 | | 2 | | 0 | | 2 | | 0 | | 3 | | 0 | | 5 |
|  | **D434** | | **N** | 0 | | 1 | | 0 | | 1 | | 1 | | 4 | | 1 | | 6 |
|  | **D229** | | **nsp1** | 0 | | 1 | | 0 | | 1 | | 0 | | 1 | | 0 | | 1 |
|  | **D276** | | **nsp1** | 0 | | 1 | | 0 | | 1 | | 0 | | 1 | | 0 | | 1 |
|  | **D277** | | **nsp1** | 0 | | 0 | | 0 | | 0 | | 0 | | 2 | | 0 | | 2 |
|  | **D285** | | **nsp1** | 0 | | 0 | | 0 | | 0 | | 0 | | 2 | | 0 | | 2 |
|  | **D290** | | **nsp1** | 0 | | 0 | | 0 | | 0 | | 0 | | 2 | | 0 | | 2 |
| **D329** | | **nsp1** | | | 0 | 0 | 0 | | 0 | | 0 | | 2 | | 0 | | 2 | |
| **D330** | | **nsp1** | | | 0 | 0 | 0 | | 0 | | 0 | | 2 | | 0 | | 2 | |
| **D378** | | **nsp1** | | | 0 | 0 | 0 | | 0 | | 0 | | 2 | | 0 | | 2 | |
| **D431** | | **nsp1** | | | 0 | 1 | 0 | | 1 | | 0 | | 2 | | 0 | | 2 | |
| **D432** | | **nsp1** | | | 0 | 0 | 0 | | 0 | | 0 | | 2 | | 0 | | 2 | |
| **D434** | | **nsp1** | | | 0 | 0 | 0 | | 0 | | 0 | | 2 | | 0 | | 2 | |
| **D229** | | **nsp11** | | | NC | NC | NC | | NC | | NC | | NC | | NC | | NC | |
| **D276** | | **nsp11** | | | NC | NC | NC | | NC | | NC | | NC | | NC | | NC | |
| **D277** | | **nsp11** | | | NC | NC | NC | | NC | | NC | | NC | | NC | | NC | |
| **D285** | | **nsp11** | | | NC | NC | NC | | NC | | NC | | NC | | NC | | NC | |
| **D330** | | **nsp11** | | | NC | NC | NC | | NC | | NC | | NC | | NC | | NC | |
| **D378** | | **nsp11** | | | NC | NC | NC | | NC | | NC | | NC | | NC | | NC | |
| **D0** | | **nsp12** | | | 0 | 0 | 0 | | 0 | | 0 | | 0 | | 0 | | 3 | |
| **D229** | | **nsp12** | | | 1 | 0 | 1 | | 0 | | 0 | | 0 | | 0 | | 3 | |
| **D276** | | **nsp12** | | | 1 | 0 | 1 | | 0 | | 0 | | 0 | | 0 | | 3 | |
| **D277** | | **nsp12** | | | 0 | 0 | 0 | | 0 | | 1 | | 0 | | 1 | | 3 | |
| **D285** | | **nsp12** | | | 0 | 0 | 0 | | 0 | | 0 | | 0 | | 0 | | 3 | |
| **D290** | | **nsp12** | | | 0 | 0 | 0 | | 0 | | 2 | | 0 | | 2 | | 3 | |
| **D329** | | **nsp12** | | | 0 | 0 | 0 | | 0 | | 2 | | 0 | | 2 | | 3 | |
| **D330** | | **nsp12** | | | 1 | 0 | 1 | | 0 | | 0 | | 0 | | 0 | | 3 | |
| **D378** | | **nsp12** | | | 0 | 0 | 0 | | 0 | | 1 | | 0 | | 1 | | 3 | |
| **D431** | | **nsp12** | | | 2 | 0 | 2 | | 0 | | 1 | | 0 | | 1 | | 3 | |
| **D432** | | **nsp12** | | | 1 | 0 | 1 | | 0 | | 1 | | 0 | | 1 | | 3 | |
| **D434** | | **nsp12** | | | 1 | 1 | 1 | | 1 | | 1 | | 0 | | 1 | | 3 | |
| **D0** | | **nsp13** | | | 0 | 5 | 0 | | 5 | | 0 | | 0 | | 0 | | 2 | |
| **D229** | | **nsp13** | | | 0 | 0 | 0 | | 0 | | 1 | | 2 | | 1 | | 4 | |
| **D276** | | **nsp13** | | | 0 | 1 | 0 | | 1 | | 1 | | 2 | | 1 | | 2 | |
| **D277** | | **nsp13** | | | 0 | 0 | 0 | | 0 | | 1 | | 3 | | 1 | | 5 | |
| **D285** | | **nsp13** | | | 0 | 0 | 0 | | 0 | | 1 | | 2 | | 1 | | 4 | |
| **D290** | | **nsp13** | | | 0 | 0 | 0 | | 0 | | 1 | | 3 | | 1 | | 5 | |
| **D329** | | **nsp13** | | | 0 | 0 | 0 | | 0 | | 1 | | 3 | | 1 | | 5 | |
| **D330** | | **nsp13** | | | 1 | 1 | 1 | | 1 | | 1 | | 2 | | 1 | | 4 | |
| **D378** | | **nsp13** | | | 0 | 0 | 0 | | 0 | | 1 | | 4 | | 1 | | 6 | |
| **D431** | | **nsp13** | | | 2 | 1 | 2 | | 1 | | 1 | | 3 | | 1 | | 5 | |
| **D432** | | **nsp13** | | | 2 | 0 | 2 | | 0 | | 1 | | 4 | | 1 | | 6 | |
| **D434** | | **nsp13** | | | 0 | 2 | 0 | | 2 | | 1 | | 3 | | 1 | | 5 | |
| **D0** | | **nsp14** | | | 0 | 0 | 0 | | 0 | | 0 | | 0 | | 1 | | 0 | |
| **D229** | | **nsp14** | | | 0 | 0 | 1 | | 0 | | 0 | | 0 | | 0 | | 0 | |
| **D276** | | **nsp14** | | | 0 | 0 | 0 | | 0 | | 0 | | 0 | | 1 | | 0 | |
| **D277** | | **nsp14** | | | 0 | 0 | 0 | | 0 | | 0 | | 0 | | 1 | | 0 | |
| **D285** | | **nsp14** | | | 0 | 0 | 1 | | 0 | | 0 | | 0 | | 0 | | 0 | |
| **D290** | | **nsp14** | | | 0 | 0 | 0 | | 0 | | 0 | | 0 | | 1 | | 0 | |
| **D329** | | **nsp14** | | | 0 | 0 | 0 | | 0 | | 0 | | 0 | | 1 | | 0 | |
| **D330** | | **nsp14** | | | 0 | 0 | 1 | | 0 | | 0 | | 0 | | 0 | | 0 | |
| **D378** | | **nsp14** | | | 0 | 0 | 0 | | 0 | | 0 | | 0 | | 1 | | 0 | |
| **D434** | | **nsp14** | | | 0 | 0 | 1 | | 0 | | 0 | | 0 | | 0 | | 0 | |
| **D378** | | **nsp15** | | | 1 | 0 | 1 | | 0 | | 0 | | 0 | | 0 | | 0 | |
| **D229** | | **nsp16** | | | 0 | 1 | 0 | | 1 | | 0 | | 0 | | 0 | | 0 | |
| **D285** | | **nsp16** | | | 0 | 1 | 0 | | 1 | | 0 | | 0 | | 0 | | 0 | |
| **D431** | | **nsp16** | | | 2 | 1 | 2 | | 1 | | 0 | | 0 | | 0 | | 0 | |
| **D432** | | **nsp16** | | | 1 | 1 | 1 | | 1 | | 1 | | 0 | | 1 | | 0 | |
| **D229** | | **nsp2** | | | 0 | 0 | 1 | | 0 | | 0 | | 1 | | 0 | | 1 | |
| **D276** | | **nsp2** | | | 0 | 0 | 1 | | 0 | | 0 | | 0 | | 0 | | 0 | |
| **D277** | | **nsp2** | | | 0 | 1 | 0 | | 1 | | 0 | | 0 | | 1 | | 0 | |
| **D285** | | **nsp2** | | | 0 | 0 | 0 | | 0 | | 0 | | 2 | | 1 | | 2 | |
| **D290** | | **nsp2** | | | 0 | 1 | 0 | | 1 | | 0 | | 0 | | 1 | | 0 | |
| **D329** | | **nsp2** | | | 0 | 1 | 0 | | 1 | | 0 | | 0 | | 1 | | 0 | |
| **D330** | | **nsp2** | | | 0 | 1 | 0 | | 1 | | 0 | | 2 | | 1 | | 2 | |
| **D378** | | **nsp2** | | | 0 | 0 | 0 | | 0 | | 2 | | 1 | | 3 | | 1 | |
| **D431** | | **nsp2** | | | 1 | 3 | 1 | | 3 | | 0 | | 4 | | 1 | | 4 | |
| **D432** | | **nsp2** | | | 1 | 3 | 1 | | 3 | | 0 | | 4 | | 1 | | 4 | |
| **D434** | | **nsp2** | | | 2 | 2 | 2 | | 2 | | 0 | | 3 | | 1 | | 3 | |
| **D0** | | **nsp3** | | | 0 | 0 | 0 | | 0 | | 0 | | 0 | | 3 | | 0 | |
| **D229** | | **nsp3** | | | 1 | 4 | 1 | | 5 | | 0 | | 3 | | 3 | | 3 | |
| **D276** | | **nsp3** | | | 1 | 0 | 1 | | 1 | | 0 | | 3 | | 2 | | 3 | |
| **D277** | | **nsp3** | | | 0 | 4 | 0 | | 4 | | 1 | | 3 | | 4 | | 4 | |
| **D285** | | **nsp3** | | | 1 | 3 | 1 | | 3 | | 1 | | 3 | | 4 | | 3 | |
| **D290** | | **nsp3** | | | 0 | 5 | 0 | | 5 | | 1 | | 4 | | 4 | | 5 | |
| **D329** | | **nsp3** | | | 1 | 0 | 2 | | 0 | | 2 | | 5 | | 5 | | 6 | |
| **D330** | | **nsp3** | | | 3 | 3 | 3 | | 4 | | 1 | | 3 | | 4 | | 3 | |
| **D378** | | **nsp3** | | | 0 | 0 | 0 | | 0 | | 1 | | 4 | | 4 | | 5 | |
| **D431** | | **nsp3** | | | 4 | 0 | 4 | | 0 | | 1 | | 6 | | 4 | | 6 | |
| **D432** | | **nsp3** | | | 3 | 0 | 3 | | 0 | | 1 | | 6 | | 4 | | 6 | |
| **D434** | | **nsp3** | | | 1 | 1 | 1 | | 2 | | 0 | | 6 | | 3 | | 6 | |
| **D0** | | **nsp4** | | | 0 | 0 | 0 | | 0 | | 0 | | 0 | | 0 | | 1 | |
| **D229** | | **nsp4** | | | 0 | 1 | 0 | | 1 | | 1 | | 3 | | 1 | | 4 | |
| **D276** | | **nsp4** | | | NC | NC | NC | | NC | | NC | | NC | | NC | | NC | |
| **D277** | | **nsp4** | | | 1 | 0 | 1 | | 0 | | 0 | | 2 | | 0 | | 3 | |
| **D285** | | **nsp4** | | | 0 | 0 | 0 | | 0 | | 1 | | 3 | | 1 | | 4 | |
| **D290** | | **nsp4** | | | 1 | 0 | 1 | | 0 | | 0 | | 3 | | 0 | | 4 | |
| **D329** | | **nsp4** | | | 0 | 0 | 0 | | 0 | | 0 | | 3 | | 0 | | 4 | |
| **D330** | | **nsp4** | | | 0 | 0 | 0 | | 0 | | 1 | | 2 | | 1 | | 3 | |
| **D378** | | **nsp4** | | | 2 | 0 | 2 | | 0 | | 0 | | 2 | | 0 | | 3 | |
| **D431** | | **nsp4** | | | 0 | 1 | 0 | | 1 | | 1 | | 3 | | 1 | | 4 | |
| **D432** | | **nsp4** | | | 0 | 1 | 0 | | 1 | | 1 | | 4 | | 1 | | 5 | |
| **D434** | | **nsp4** | | | 0 | 0 | 0 | | 0 | | 1 | | 3 | | 1 | | 4 | |
| **D229** | | **nsp5** | | | 0 | 1 | 0 | | 1 | | 0 | | 0 | | 1 | | 4 | |
| **D378** | | **nsp5** | | | 1 | 0 | 1 | | 0 | | 0 | | 0 | | 0 | | 0 | |
| **D434** | | **nsp5** | | | 1 | 0 | 1 | | 0 | | 0 | | 0 | | 0 | | 0 | |
| **D276** | | **nsp5** | | | 0 | 0 | 0 | | 0 | | 0 | | 0 | | 1 | | 2 | |
| **D290** | | **nsp5** | | | 1 | 0 | 1 | | 0 | | 0 | | 0 | | 0 | | 0 | |
| **D285** | | **nsp5** | | | 0 | 0 | 0 | | 0 | | 0 | | 0 | | 0 | | 0 | |
| **D431** | | **nsp5** | | | 0 | 0 | 0 | | 1 | | 0 | | 0 | | 0 | | 0 | |
| **D432** | | **nsp5** | | | 0 | 1 | 0 | | 0 | | 0 | | 0 | | 0 | | 0 | |
| **D277** | | **nsp5** | | | 1 | 0 | 1 | | 0 | | 0 | | 0 | | 0 | | 3 | |
| **D0** | | **nsp5** | | | 0 | 0 | 0 | | 0 | | 0 | | 0 | | 0 | | 1 | |
| **D0** | | **nsp6** | | | 0 | 0 | 0 | | 0 | | 0 | | 0 | | 1 | | 0 | |
| **D229** | | **nsp6** | | | 0 | 0 | 0 | | 0 | | 0 | | 1 | | 1 | | 1 | |
| **D276** | | **nsp6** | | | 0 | 0 | 0 | | 0 | | 0 | | 1 | | 0 | | 1 | |
| **D277** | | **nsp6** | | | 0 | 0 | 0 | | 0 | | 0 | | 1 | | 1 | | 1 | |
| **D285** | | **nsp6** | | | 0 | 0 | 0 | | 0 | | 0 | | 1 | | 1 | | 1 | |
| **D290** | | **nsp6** | | | 0 | 0 | 0 | | 0 | | 0 | | 1 | | 1 | | 1 | |
| **D329** | | **nsp6** | | | 0 | 0 | 0 | | 0 | | 0 | | 1 | | 1 | | 1 | |
| **D330** | | **nsp6** | | | 1 | 0 | 1 | | 0 | | 0 | | 1 | | 1 | | 1 | |
| **D378** | | **nsp6** | | | 0 | 0 | 0 | | 0 | | 0 | | 1 | | 1 | | 1 | |
| **D431** | | **nsp6** | | | 0 | 0 | 0 | | 0 | | 1 | | 1 | | 2 | | 1 | |
| **D432** | | **nsp6** | | | 0 | 0 | 0 | | 0 | | 1 | | 1 | | 2 | | 1 | |
| **D434** | | **nsp6** | | | 0 | 0 | 0 | | 0 | | 1 | | 1 | | 2 | | 1 | |
| **D276** | | **nsp8** | | | 0 | 0 | 0 | | 0 | | 1 | | 1 | | 1 | | 1 | |
| **D277** | | **nsp8** | | | 1 | 1 | 1 | | 1 | | 0 | | 0 | | 0 | | 0 | |
| **D290** | | **nsp8** | | | 1 | 1 | 1 | | 1 | | 0 | | 0 | | 0 | | 0 | |
| **D285** | | **nsp9** | | | 1 | 0 | 1 | | 0 | | 0 | | 0 | | 0 | | 0 | |
| **D329** | | **nsp9** | | | 0 | 1 | 0 | | 1 | | 0 | | 0 | | 0 | | 0 | |
| **D0** | | **ORF3a** | | | 0 | 0 | 0 | | 0 | | 0 | | 1 | | 1 | | 1 | |
| **D229** | | **ORF3a** | | | 1 | 2 | 2 | | 2 | | 0 | | 1 | | 0 | | 1 | |
| **D276** | | **ORF3a** | | | 1 | 0 | 1 | | 0 | | 0 | | 2 | | 1 | | 2 | |
| **D277** | | **ORF3a** | | | 0 | 3 | 0 | | 3 | | 1 | | 0 | | 1 | | 0 | |
| **D285** | | **ORF3a** | | | 0 | 3 | 1 | | 3 | | 0 | | 0 | | 0 | | 0 | |
| **D290** | | **ORF3a** | | | 0 | 3 | 1 | | 3 | | 1 | | 0 | | 1 | | 0 | |
| **D329** | | **ORF3a** | | | 0 | 0 | 0 | | 0 | | 0 | | 3 | | 1 | | 3 | |
| **D330** | | **ORF3a** | | | 2 | 0 | 2 | | 0 | | 0 | | 0 | | 0 | | 0 | |
| **D378** | | **ORF3a** | | | 0 | 0 | 0 | | 0 | | 1 | | 0 | | 1 | | 0 | |
| **D431** | | **ORF3a** | | | 1 | 1 | 1 | | 1 | | 0 | | 0 | | 0 | | 0 | |
| **D432** | | **ORF3a** | | | 1 | 0 | 1 | | 0 | | 0 | | 1 | | 0 | | 1 | |
| **D434** | | **ORF3a** | | | 1 | 0 | 1 | | 0 | | 0 | | 0 | | 0 | | 0 | |
| **D378** | | **ORF6** | | | 0 | 1 | 0 | | 1 | | 0 | | 0 | | 0 | | 0 | |
| **D434** | | **ORF6** | | | 0 | 1 | 0 | | 1 | | 0 | | 0 | | 0 | | 0 | |
| **D229** | | **ORF7a** | | | 0 | 1 | 0 | | 1 | | 0 | | 0 | | 0 | | 0 | |
| **D276** | | **ORF7a** | | | NC | NC | NC | | NC | | NC | | NC | | NC | | NC | |
| **D277** | | **ORF7a** | | | NC | NC | NC | | NC | | NC | | NC | | NC | | NC | |
| **D285** | | **ORF7a** | | | NC | NC | NC | | NC | | NC | | NC | | NC | | NC | |
| **D290** | | **ORF7a** | | | NC | NC | NC | | NC | | NC | | NC | | NC | | NC | |
| **D330** | | **ORF7a** | | | NC | NC | NC | | NC | | NC | | NC | | NC | | NC | |
| **D378** | | **ORF7a** | | | NC | NC | NC | | NC | | NC | | NC | | NC | | NC | |
| **D431** | | **ORF7a** | | | NC | NC | NC | | NC | | NC | | NC | | NC | | NC | |
| **D432** | | **ORF7a** | | | 0 | 0 | 0 | | 0 | | 0 | | 1 | | 0 | | 1 | |
| **D434** | | **ORF7a** | | | NC | NC | NC | | NC | | NC | | NC | | NC | | NC | |
| **D229** | | **ORF8** | | | 1 | 0 | 1 | | 3 | | 0 | | 0 | | 0 | | 0 | |
| **D277** | | **ORF8** | | | 0 | 2 | 0 | | 2 | | 0 | | 0 | | 0 | | 0 | |
| **D285** | | **ORF8** | | | 0 | 2 | 0 | | 2 | | 0 | | 0 | | 0 | | 0 | |
| **D290** | | **ORF8** | | | 0 | 2 | 0 | | 2 | | 0 | | 0 | | 0 | | 0 | |
| **D229** | | **ORF9b** | | | 0 | 0 | 0 | | 0 | | 0 | | 1 | | 0 | | 1 | |
| **D276** | | **ORF9b** | | | 0 | 0 | 0 | | 0 | | 0 | | 1 | | 0 | | 1 | |
| **D277** | | **ORF9b** | | | 0 | 0 | 0 | | 0 | | 0 | | 1 | | 0 | | 1 | |
| **D285** | | **ORF9b** | | | 0 | 0 | 0 | | 0 | | 0 | | 1 | | 0 | | 1 | |
| **D290** | | **ORF9b** | | | 0 | 0 | 0 | | 0 | | 0 | | 1 | | 0 | | 1 | |
| **D329** | | **ORF9b** | | | 0 | 0 | 0 | | 0 | | 0 | | 1 | | 0 | | 1 | |
| **D330** | | **ORF9b** | | | 0 | 0 | 0 | | 0 | | 0 | | 1 | | 0 | | 1 | |
| **D378** | | **ORF9b** | | | 0 | 0 | 0 | | 0 | | 0 | | 1 | | 0 | | 1 | |
| **D431** | | **ORF9b** | | | 0 | 1 | 0 | | 1 | | 0 | | 2 | | 0 | | 2 | |
| **D432** | | **ORF9b** | | | 0 | 1 | 0 | | 1 | | 0 | | 2 | | 0 | | 2 | |
| **D434** | | **ORF9b** | | | 0 | 1 | 0 | | 1 | | 0 | | 3 | | 0 | | 3 | |
| **D0** | | **S** | | | 1 | 0 | 1 | | 0 | | 0 | | 0 | | 0 | | 2 | |
| **D229** | | **S** | | | 1 | 2 | 1 | | 7 | | 1 | | 1 | | 1 | | 2 | |
| **D276** | | **S** | | | 0 | 1 | 0 | | 6 | | 1 | | 2 | | 1 | | 3 | |
| **D277** | | **S** | | | 1 | 4 | 1 | | 4 | | 0 | | 0 | | 0 | | 6 | |
| **D285** | | **S** | | | 2 | 5 | 2 | | 6 | | 0 | | 0 | | 0 | | 7 | |
| **D290** | | **S** | | | 1 | 3 | 1 | | 4 | | 0 | | 0 | | 0 | | 7 | |
| **D329** | | **S** | | | 0 | 1 | 0 | | 1 | | 0 | | 1 | | 0 | | 8 | |
| **D330** | | **S** | | | 0 | 2 | 0 | | 2 | | 0 | | 0 | | 0 | | 6 | |
| **D378** | | **S** | | | 0 | 1 | 0 | | 1 | | 0 | | 0 | | 0 | | 6 | |
| **D431** | | **S** | | | 0 | 0 | 0 | | 0 | | 0 | | 1 | | 0 | | 8 | |
| **D432** | | **S** | | | 0 | 0 | 0 | | 0 | | 0 | | 1 | | 0 | | 8 | |
| **D434** | | **S** | | | 0 | 2 | 0 | | 2 | | 0 | | 1 | | 0 | | 8 | |

Table S2

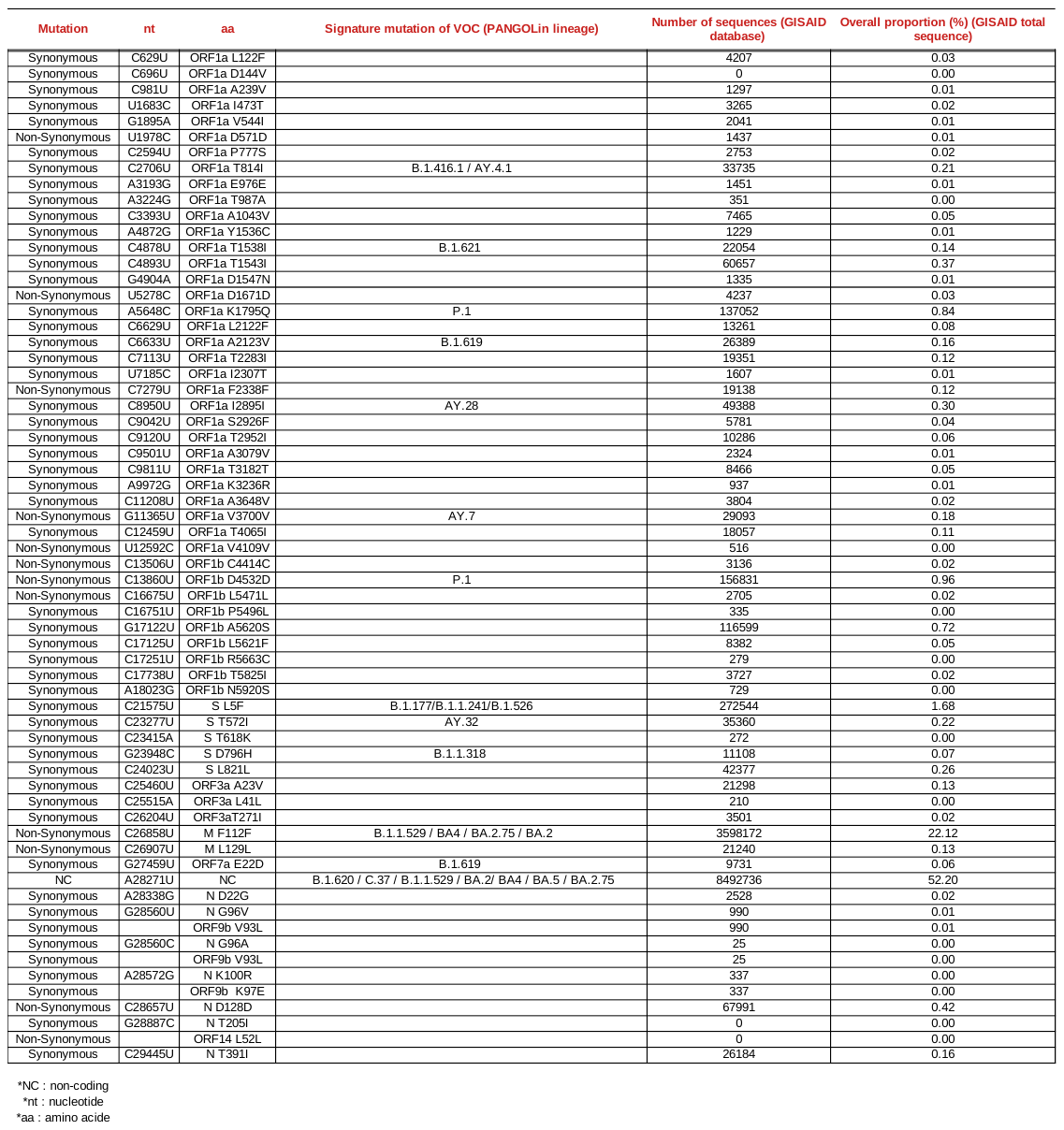

Table S3

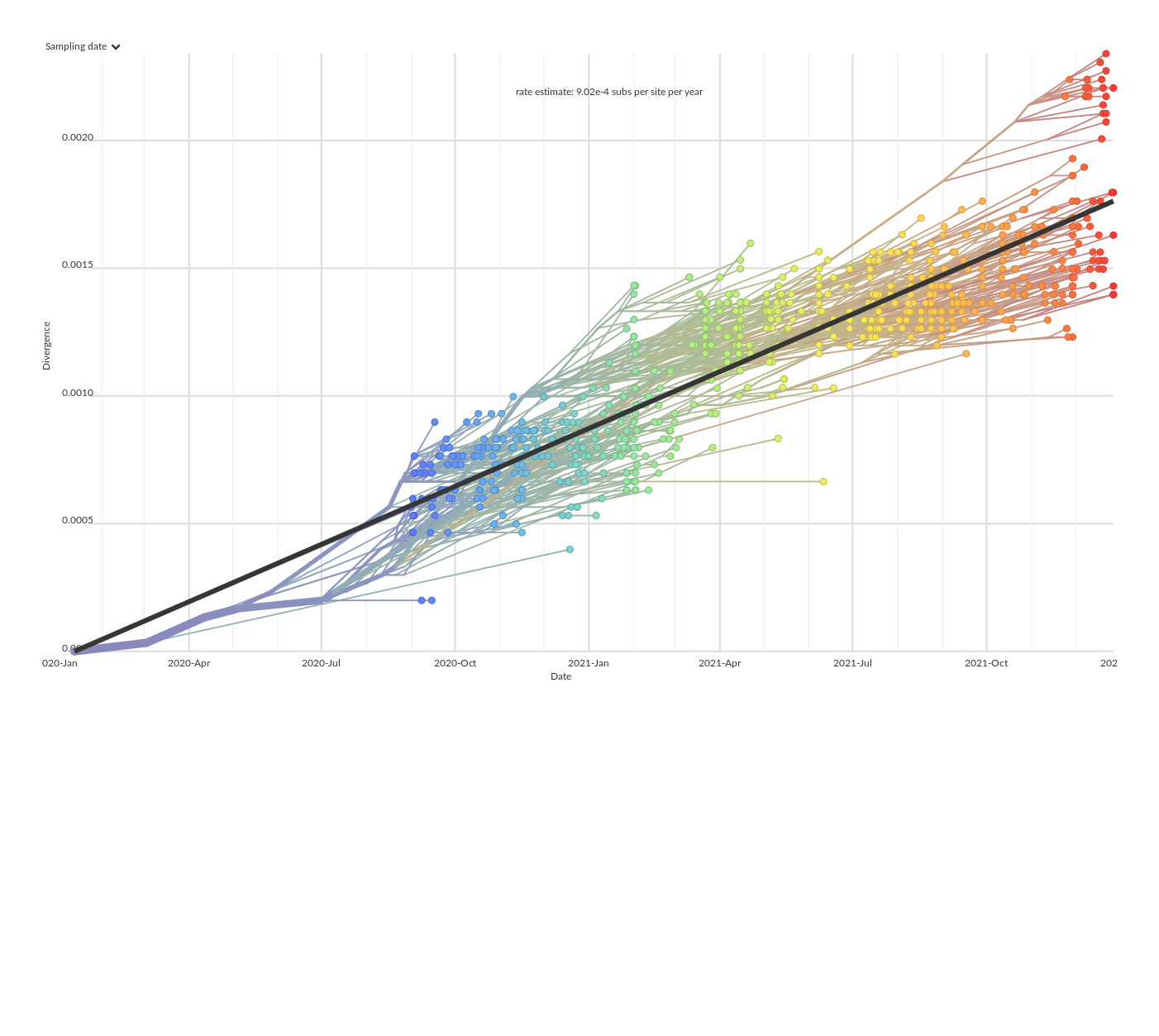
FIGURES

Figure S1

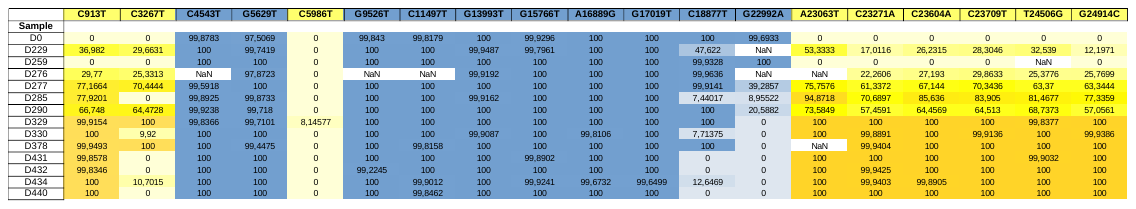

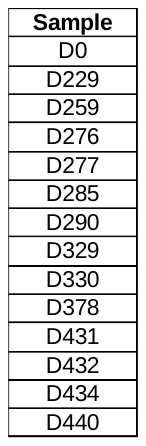

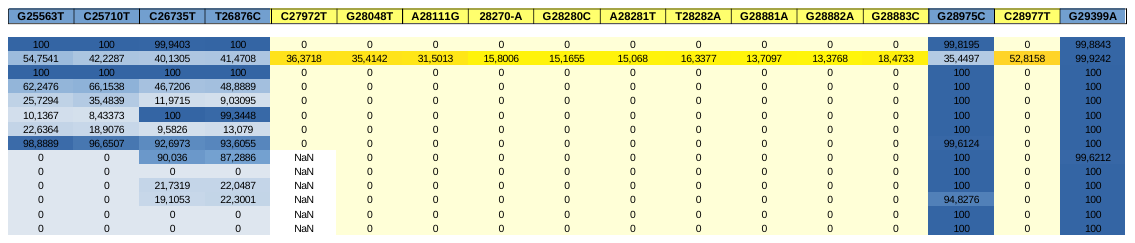

Figure S2

Figure S3

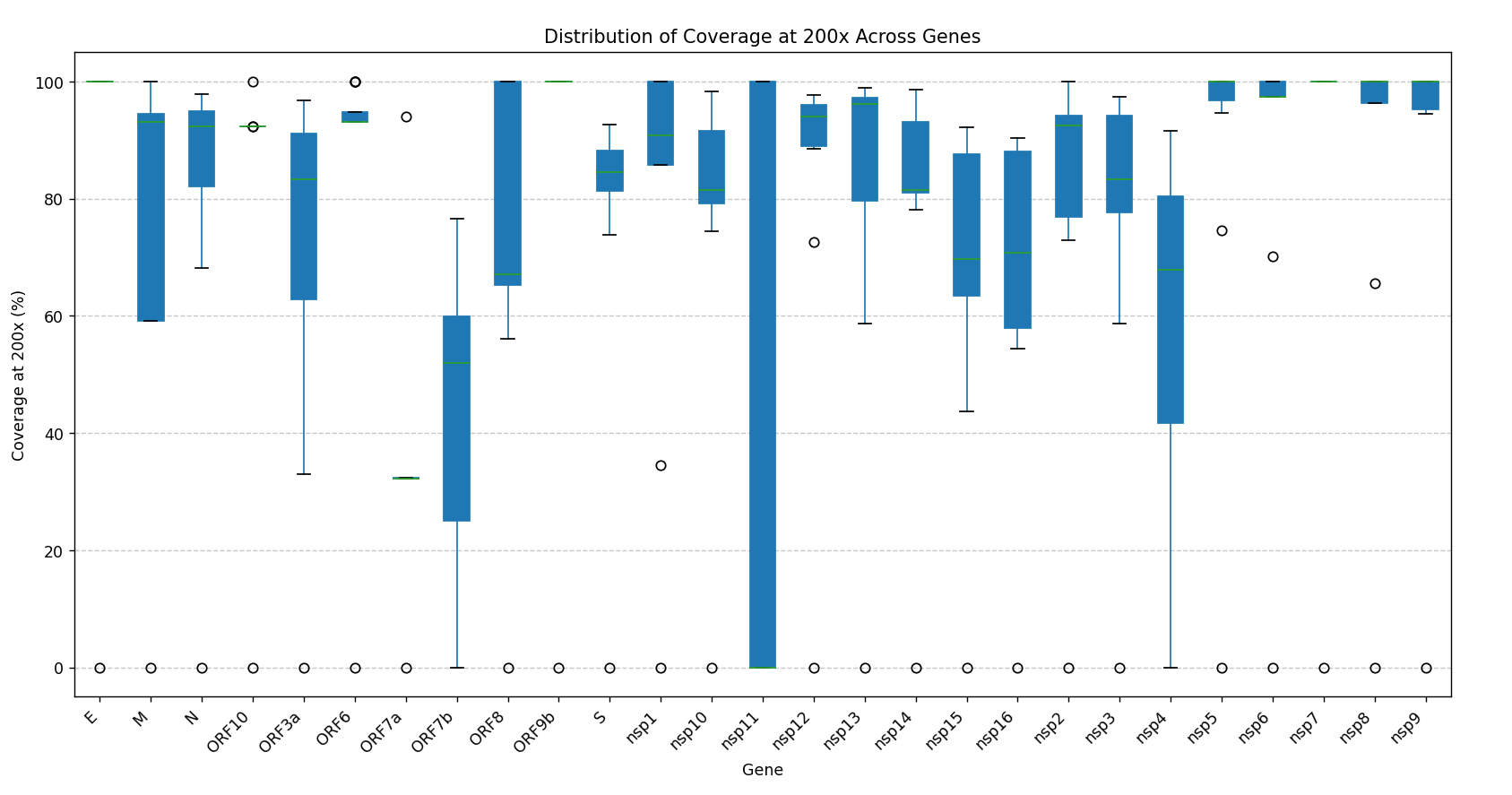

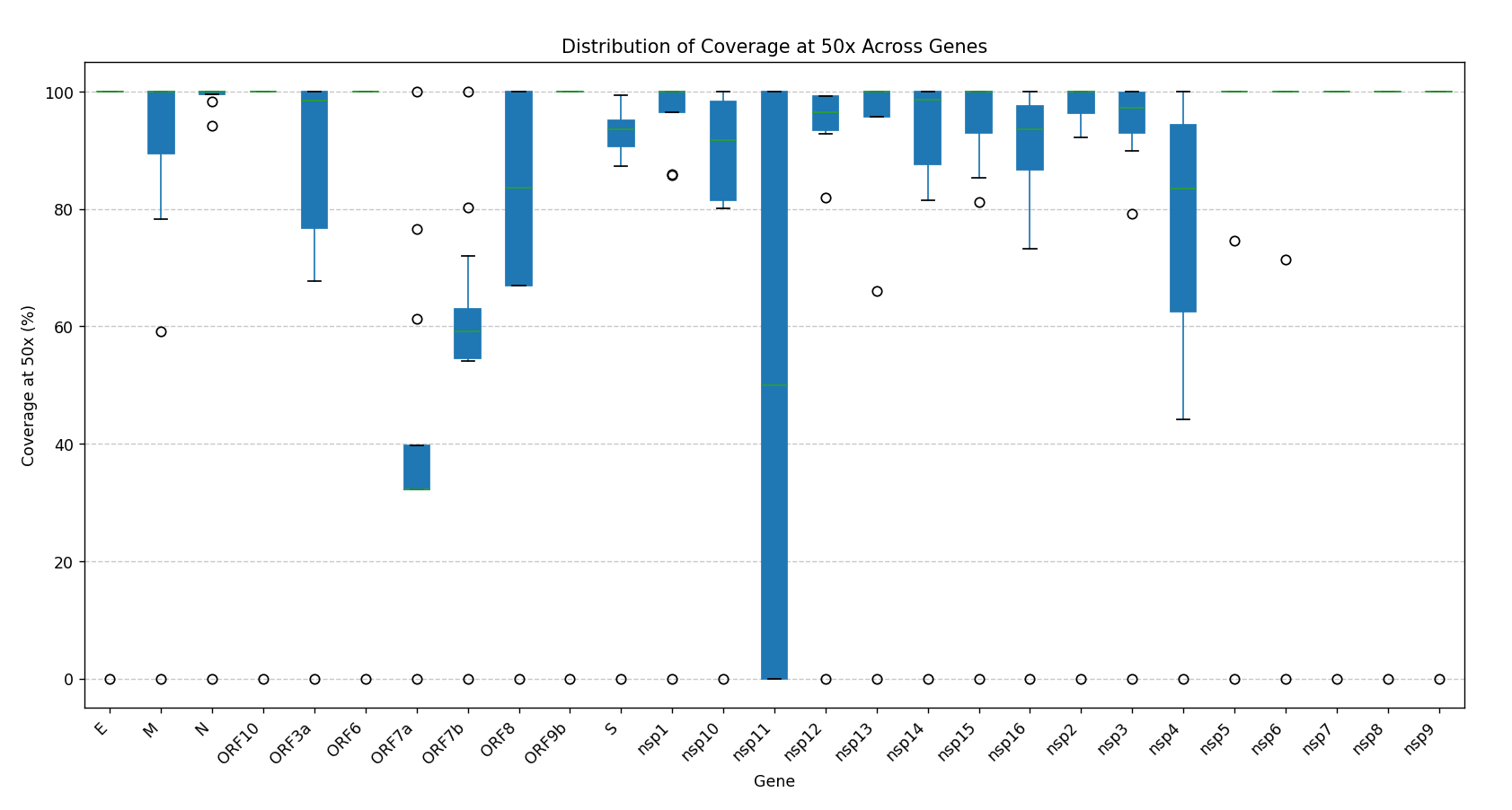

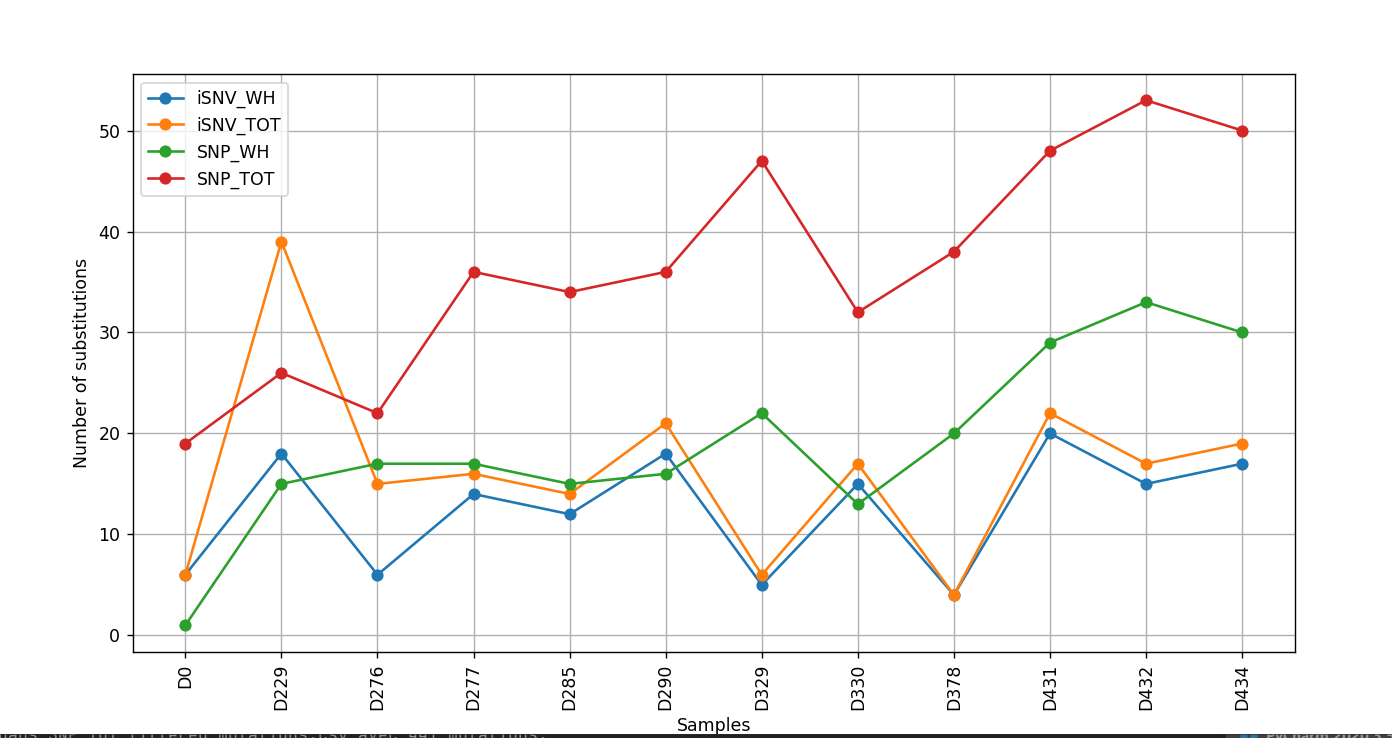

Cov. 200x

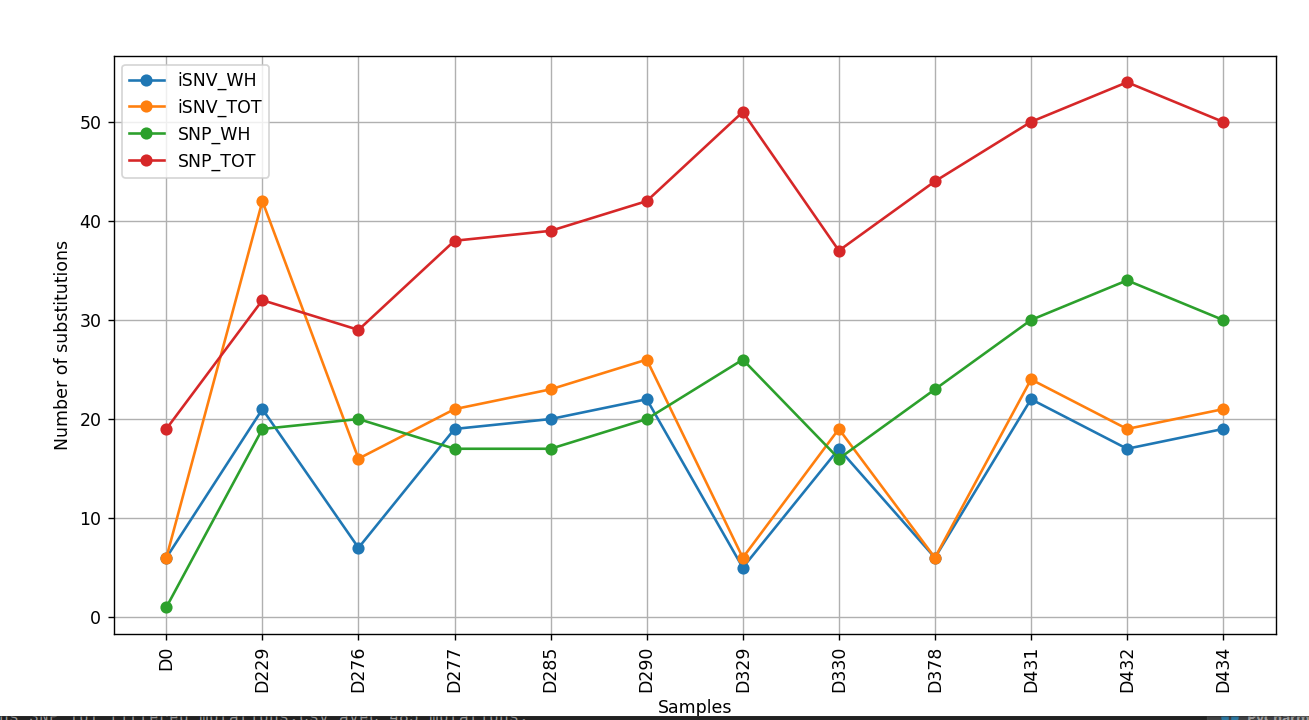

Cov. 50x

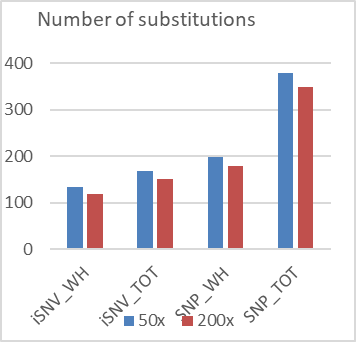

a)

b)

Cov. 50x

Cov. 200x

c)

d)

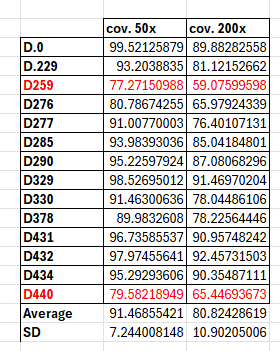

Number of substitutions

|  | 50x | 200x |
| --- | --- | --- |
| iSNV_WH | 135 | 118 |
| iSNV_TOT | 169 | 151 |
| SNP_WH | 199 | 179 |
| SNP_TOT | 379 | 349 |

Figure S3

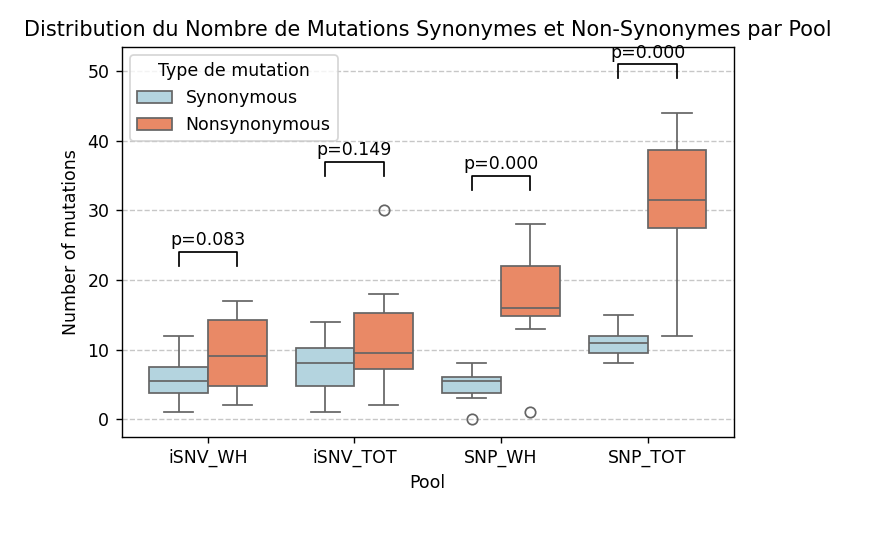

Figure S4

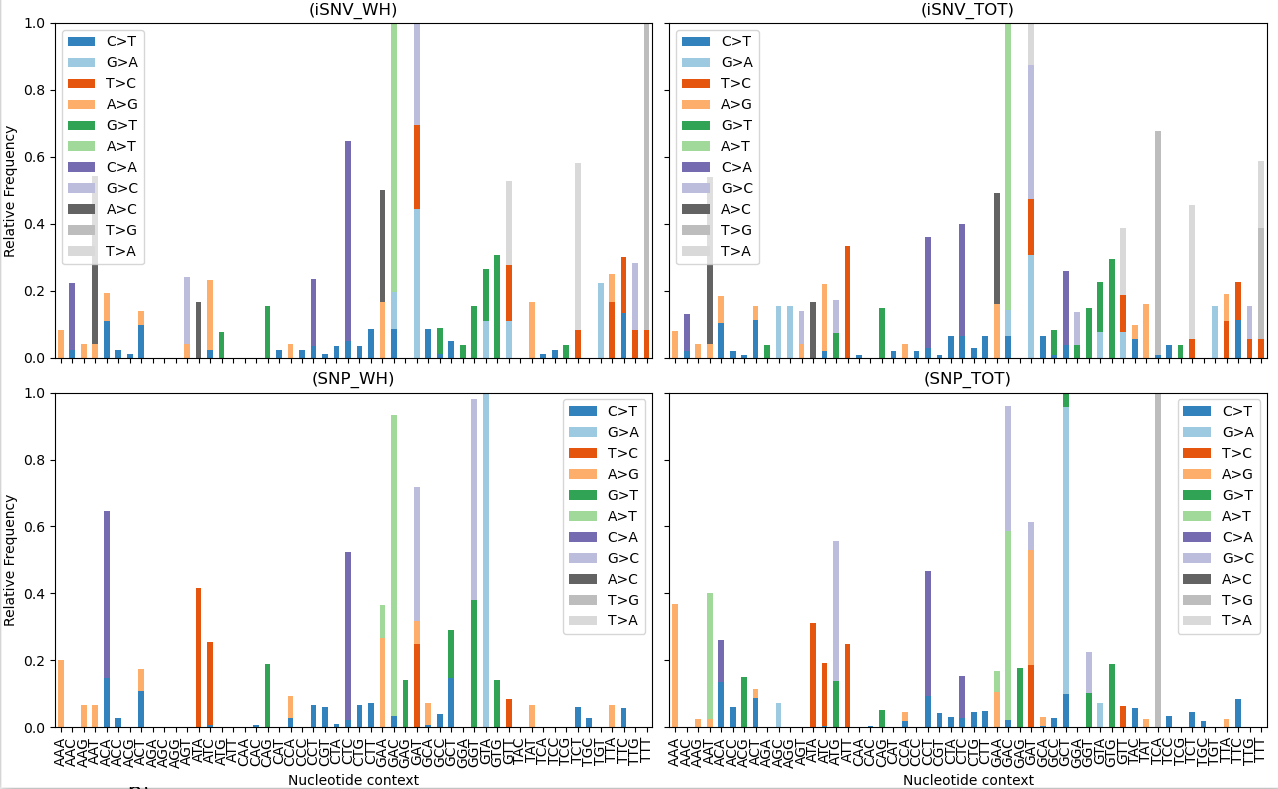

Figure S5

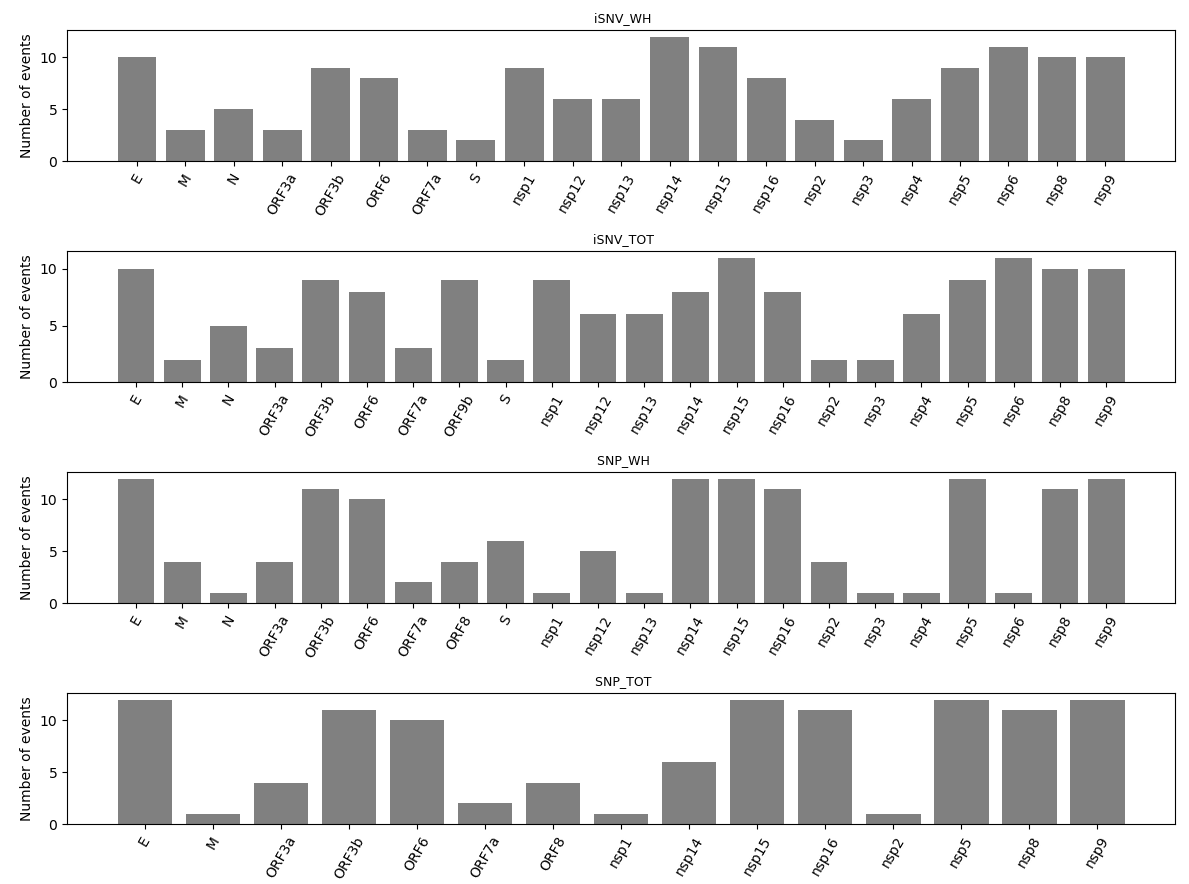

Figure S6 : Indeterminate value : pN=0 and pS=0

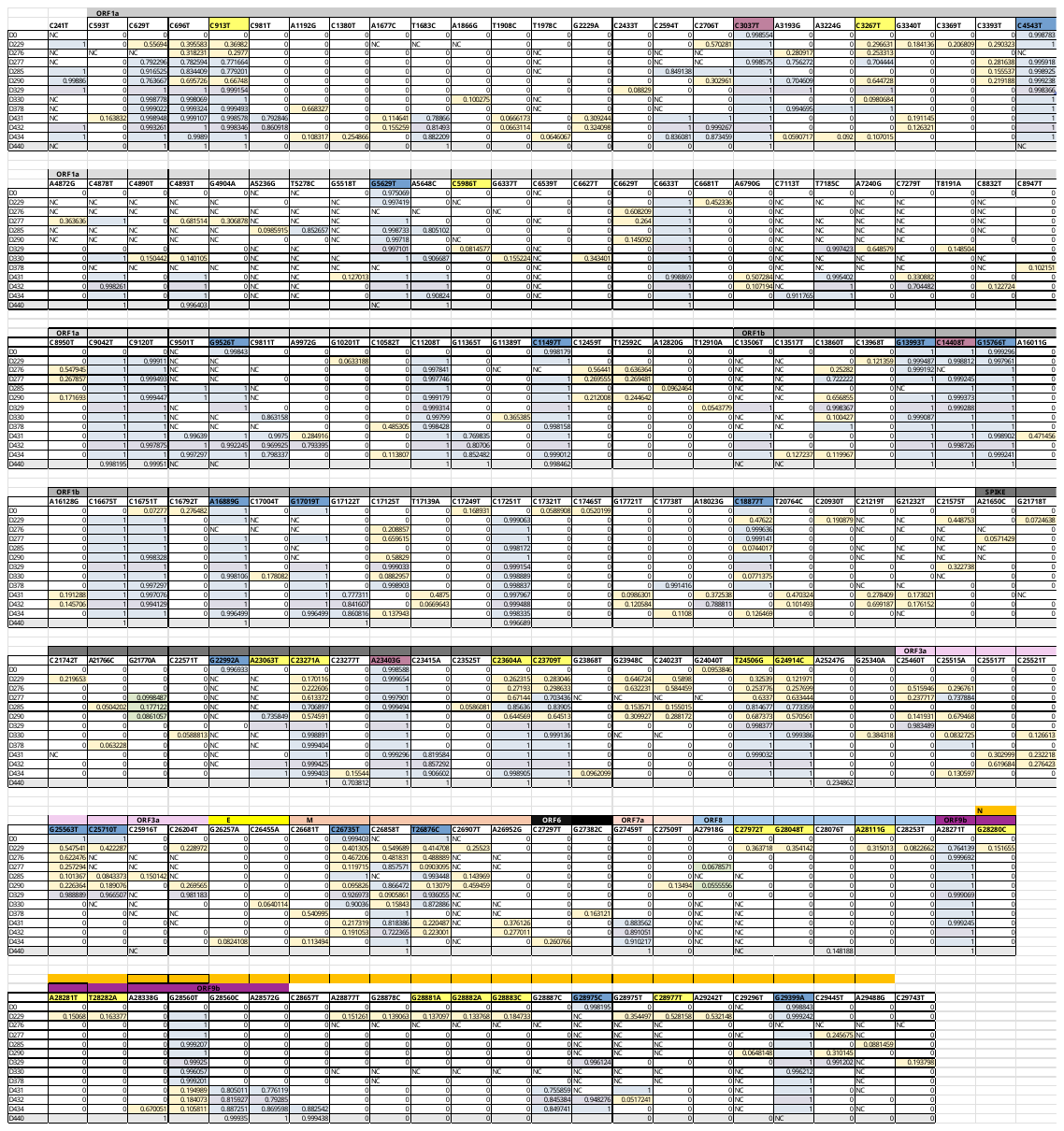

Figure S7 Mutations associated with allele frequency

LEGEND OF SUPPLEMENTARY

Table S1

Nextclade nomenclature (column 1) showing the different genotypes of B.1.1.7 circulating in Marseille from January 2020 to May 2021. We noticed the circulation of seven different B.1.1.7 genotypes. No classified as “Within host” mutation was found between them.

Table S2

Table of non-synonymous and synonymous mutation count within each pool of mutations. Genes are indicated in column 2.

Figure S1

Time-scaled maximum-likelihood phylogenetic tree constructed using Nextstrain©, comprising 700 sequences randomly selected locally in the GISAID database to cover the period from January 2020 to December 2021. Time is represented on x-axis and divergence on the y-axis.

Figure S2

Figure Legend: Table representing the intra-host sequential frequencies of mutations resulting exclusively from recombination events between parental genomes B.1.160 and B.1.1.7. Sampling dates are indicated by D0, D229, D259, etc., representing different time points during the intra-host evolution. Mutations highlighted in blue correspond to those associated with the B.1.160 variant, while mutations highlighted in yellow represent those from the B.1.1.7 variant. The frequencies (%) of each mutation at each time point are shown in the cells. Cells marked as NaN indicate regions not covered by sequencing or missing data.

Figure S3

a) Table indicating coverage (% of genome covered) at 50 and 200 reads depth. In red, genomes with < 80% coverage. b) Percentage of genes covered at 50 (top panel) and 200 (bottom panel) reads depth. c) Number of mutations counted in each pool for the 12 samples selected, at 50 and 200 read depth.

Figure S4

Boxplot comparing the number of synonymous and non-synonymous mutations in the four pools, with statistical significance indicated by p-values (Wilcoxon test).

Figure S5

Relative frequency of the different mutation types associated with their nucleotide contexts (x-axis), meaning the percentage of mutation type within a trinucleotide context.

Figure S6

Histogram showing the number of samples for which the ratio could not be calculated with pN=0 and pS=0, leading to indeterminate selection.

Figure S7

The parental genomes are highlighted with blue (B.1.160), yellow (B.1.1.7),purple for ancestral mutation, NC meaning “Not covered”,
